# Polymorphic 3D genome architecture mediated by transposable elements

**DOI:** 10.64898/2026.01.19.697799

**Authors:** Harsh Shukla, Yuheng Huang, Zita Y. Gao, Yuh Chwen G. Lee

## Abstract

The three-dimensional (3D) folding of the genome plays a crucial role in genome regulation. However, how 3D genome structure varies between individuals and consequently influences genome function and evolution remains poorly understood. One potential source of this variation is transposable elements (TEs), genomic parasites whose location and composition vary between species and individuals. Hosts typically silence TEs through enriching them with repressive epigenetic marks, turning euchromatic TEs into heterochromatin islands, which were shown to spatially interact with pericentromeric heterochromatin (PCH). Because most TE insertions are present in only a few individuals within Drosophila populations, we asked whether polymorphism in the presence/absence of TEs drives varying 3D structures through TE-PCH spatial interactions. We performed deep-coverage Hi-C of two wild-type Drosophila strains and developed a Hi-C analysis framework enabling allelic comparisons of spatial interactions with PCH. Supporting our hypothesis, nearly 40% of strain-specific euchromatic TEs cause their adjacent euchromatic regions to be spatially closer to PCH than TE-free homologous alleles. These interactions are not limited to specific TE families, and, surprisingly, telomere-proximal TEs show a similar propensity as centromere-proximal TEs to enhance PCH interactions. The most defining feature of TEs involved in PCH interactions is H3K9me3 enrichment, revealing a chromatin-based mechanism for TE-mediated 3D genome organization broadly applicable across TE families and genome locations. Importantly, TEs involved in PCH interactions reduce the expression of adjacent genes and are evolutionarily young, indicating stronger selection against them. Our study reveals a previously uncharacterized mechanism by which TEs influence the function and evolution of host genomes by generating polymorphic 3D genome organization.

## Introduction

The spatial packaging of linear genome into a nucleus several orders of magnitude smaller is not merely an effective solution to accommodate a large amount of genetic materials, but also a critical regulatory mechanism for various genome functions, such as gene expression (Bystricky and Merkenschlager 2020; Li et al. 2018; Ibrahim and Mundlos 2020), DNA replication (Ryba et al. 2010; Klein et al. 2021), and chromosome segregation (Nagano et al. 2017; Naumova et al. 2013; Zhang and Blobel 2023) [reviewed in (Giles et al. 2025; Dekker and Mirny 2024)]. Disruption of this spatial packaging, or 3D genome organization, can result in multiple disease phenotypes (Spielmann et al. 2018; Norton and Phillips-Cremins 2017). Yet, despite these critical regulatory roles, 3D genome organization is not invariant and has diverged between species (Hoencamp et al. 2021; Torosin et al. 2020, 2022), leading to functional consequences including altered gene expression (Eres et al. 2019; Wu et al. 2025). Because between-species divergence must have originated as polymorphisms within ancestral populations, characterizing intraspecies variation in 3D genome architecture is essential for understanding how nuclear organization evolves. Although naturally occurring variation in 3D genome structure is increasingly being documented within species (Gilbertson et al. 2024; McArthur et al. 2025; Li et al. 2024), the molecular and genetic drivers underlying this variation remain largely unexplored.

Identifying the drivers of varying 3D genome organization requires understanding the mechanisms that establish genome spatial structure in the first place. A fundamental feature of this spatial organization is the compartmentalization of the genome into the gene-rich, transcriptionally active euchromatin and the gene-poor, repeat-rich, and transcriptionally inert heterochromatin (reviewed in (Dekker and Mirny 2024)). This non-random spatial partitioning reflects the distinct chromatin states that regulate genome function and maintain boundaries between active and inactive genomic regions. Both experimental manipulation (Stutzman et al. 2024; Zenk et al. 2021; Falk et al. 2019) and polymer simulations (Falk et al. 2019) demonstrate that the self-associations of heterochromatin from different chromosomes primarily drive this compartmentalization. These self-associations of heterochromatin are mediated by the distinctive epigenetic modification of heterochromatin (Janssen et al. 2018; Penagos-Puig and Furlan-Magaril 2020), mainly di- and trimethylation of histone H3 at lysine 9 (H3K9me2/3). In turn, H3K9me2/3 recruit Heterochromatin Protein 1a (HP1a), a key architectural protein of heterochromatin (James and Elgin 1986), whose phase separation properties drive heterochromatin coalescence (Strom et al. 2017; Larson et al. 2017; Keenen et al. 2021).

Interestingly, the enrichment of H3K9me2/3 and HP1a is not restricted to constitutive heterochromatin, but also occurs at discrete locations throughout gene-dense euchromatin (Riddle et al. 2011; Becker et al. 2017), with a substantial fraction of them generated by transposable elements (TEs) in *Drosophila* (Lee 2015; Lee and Karpen 2017; Huang et al. 2022). TEs are selfish genetic elements that occupy appreciable proportions of nearly all eukaryotic genomes (Wells and Feschotte 2020). Their replicative movement and mere presence can impair genome function through multiple mechanisms, including inserting into and disrupting genic sequences (Bellen et al. 2004; Maksakova et al. 2006), acting as ectopic regulatory sequences (Chuong et al. 2017), and mediating illegitimate nonhomologous recombination (Symer et al. 2002; Langley et al. 1988; Montgomery et al. 1987). To mitigate the harmful effects of TEs within euchromatin, eukaryotes commonly employ small RNA-directed enrichment of repressive marks, predominantly H3K9me2/3, at TEs (reviewed in (Slotkin and Martienssen 2007; Czech et al. 2018)). Moreover, these repressive marks frequently “spread” into neighboring euchromatic regions, converting TEs and their adjacent regions into heterochromatic islands within euchromatin ((Ahmed et al. 2011; Quadrana et al. 2016; Stuart et al. 2016; Rebollo et al. 2011; Eichten et al. 2012), reviewed in (Choi and Lee 2020)). Because these TE-mediated heterochromatin islands share molecular features with constitutive heterochromatin, we previously hypothesized and demonstrated that silenced euchromatic TEs spatially interact with constitutive heterochromatin via phase-separation mechanisms similar to those driving heterochromatin coalescence (Lee et al. 2020). These findings revealed an unexpected architectural role for TEs in shaping 3D genome organization through mediating interactions across supposedly spatially segregated genomic compartments.

However, TEs exhibit substantial differences in abundance, composition, and insertion locations across species (Wells and Feschotte 2020; Osmanski et al. 2023; Elliott and Gregory 2015), predicting that TE-mediated 3D genome organization should be highly variable. Moreover, TEs can differ substantially between individuals within the same species, particularly in their insertion locations. For example, the insertion locations of TEs vary greatly between human individuals (Sudmant et al. 2015; Stewart et al. 2011), such that any two unrelated humans differ by at least a thousand presence/absence polymorphisms of TE insertions (Bourque et al. 2018). Within *Drosophila*, TE insertions exhibit a highly skewed frequency distribution, with the majority present in only a few individuals within a population (Cridland et al. 2013; Rech et al. 2022). These presence/absence polymorphisms of TEs generate between-strain variation in H3K9me2/3 enrichment, resulting in segregating heterochromatic islands within euchromatin (Choi and Lee 2020; Huang et al. 2022). Given the prevalence of TEs and their substantial variability in insertion location, spatial interactions between polymorphic euchromatic TEs and constitutive heterochromatin could represent a powerful source of individual variation in 3D genome organization.

To investigate our hypothesis that polymorphic TE insertions drive between-individual differences in 3D genome architecture on a genome-wide scale, we generated deep-coverage High-throughput Chromosome Conformation Capture (Hi-C) data for two wildtype inbred strains of the Drosophila Synthetic Population Resource (DSPR), which were collected from distinct geographic locations (King et al. 2012). We used PacBio HiFi long-read sequencing (Wenger et al. 2019) to generate high-quality, chromosome-level genome assemblies for the studied strains, enabling precise annotation of TEs and full sequence characterization of the non-satellite portion of the highly repetitive constitutive heterochromatin. We developed a Hi-C analysis framework that, unlike previous approaches, incorporates Hi-C reads from highly repetitive heterochromatin and enables genome-wide comparisons between homologous alleles with and without TE insertions. Using this framework, we systematically investigated the contribution of TEs to varying local 3D genome structures, identified key biological properties of these TEs, and revealed their functional and evolutionary consequences. Our findings establish that polymorphic TEs act as potent architects of 3D genome variation within natural populations, revealing a previously unappreciated mechanism linking the evolutionary dynamics of TEs to the evolution of nuclear organization.

## Results

### Approaches for overcoming technical challenges in detecting 3D interactions between TEs and pericentromeric heterochromatin

We aim to investigate whether polymorphic TEs drive varying 3D structures through their spatial interactions with constitutive heterochromatin, which is usually located around the centromere and commonly referred to as pericentromeric heterochromatin, or PCH hereafter. To compare the local 3D structures of homologous sequences with and without TEs genome-wide, we conducted Hi-C (Lieberman-Aiden et al. 2009) using 16-18 hour embryos of A4 and A7 *D. melanogaster* strains from DSPR (King et al. 2012). These two inbred, wildtype strains were collected from Zimbabwe (Africa) and Taiwan (Asia), respectively, and have distinct TE insertion profiles (Chakraborty et al. 2019). Quantifying 3D interactions involving PCH poses several challenges. First, the compacted organization of PCH makes it less accessible to enzymes, leading to underrepresentation of such regions in various enzyme-based Hi-C studies (Chandradoss et al. 2020). In addition, due to the highly repetitive sequence composition of PCH, Hi-C reads coming from PCH regions are typically excluded from commonly adopted Hi-C analysis pipelines (e.g., (Sexton et al. 2012)). Relatedly, Hi-C data analysis involves mapping short reads to pre-existing genome assemblies, typically the reference genome. Such an approach often introduces reference mapping biases (Lin et al. 2024b; Ballouz et al. 2019), while genomic analysis using genome assemblies of the same strain from which the functional data were generated results in improved mapping quality (Chen et al. 2021; Shukla et al. 2019; Thorburn et al. 2023). Yet, despite the substantial genetic variation within PCH (Shukla et al. 2025; Langley et al. 2019), conducting sequence analysis using PCH genomes from the experimental strains is typically unfeasible due to challenges associated with assembling PCH (e.g., (Stutzman et al. 2024; Lee et al. 2020)).

To address these challenges, we constructed Hi-C libraries using a cocktail of four restriction enzymes with frequent cutting sites rather than a single enzyme, which enhances genome-wide fragment coverage and reduces sequence-specific biases (see Materials and Methods). In addition, we developed a Hi-C analysis pipeline that incorporates repetitive Hi-C reads and normalizes between genotypes to enable between-strains comparisons (see below). Finally, we leveraged recent advances in long-read sequencing and used Pacific Biosciences (PacBio) HiFi sequencing (Wenger et al. 2019) to construct a *de novo* assembly of one of the two strains while using the recently published HiFi assembly of the other strain (Shukla et al. 2025). Compared to the gold standard reference genome assembly of *D. melanogaster* (Hoskins et al. 2015), the two HiFi assemblies have better contiguity (Figure 1A; see Figure S1 for dotplots) and significantly improved recovery of PCH, particularly the X PCH (recovering 39.1% (A4) and 35.4% (A7) more PCH than the reference assembly; Figure 1B; PCH was defined based on strong H3K9me3 enrichment, as shown in Figure S2). These assemblies provided two advantages: they mitigate reference biases by matching the strains used in the Hi-C experiment, and their more complete PCH representation improves the detection of PCH contacts by increasing the mappable target size (see below).

**Figure 1.**
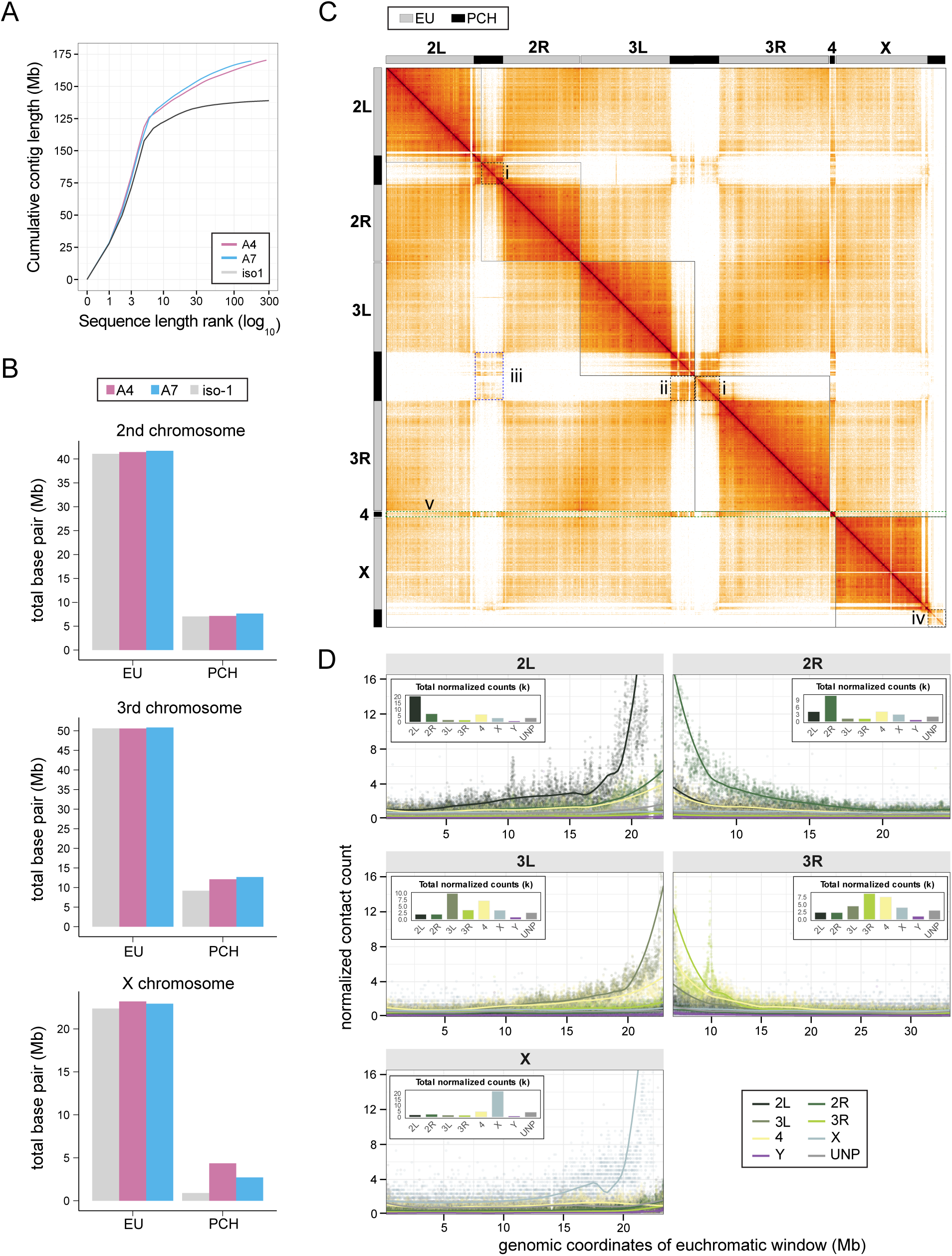
Reference-quality genome assemblies facilitate Hi-C analysis that revealed previously uncharacterized 3D interactions involving Pericentromeric Heterochromatin (PCH). **(A)** Cumulative contig length of the *de novo* PacBio HiFi assemblies for studied strains (A4: pink, A7: blue) and the *D. melanogaster* reference genome (iso-1: gray). The steeper initial slope of the studied strains relative to the reference genome indicates greater continuity in the genome assemblies of the studied strains. (**B**) Size of PCH regions of the Hi-Fi assemblies of the studied strains and the reference genome. The HiFi assemblies recover substantially more PCH sequence compared to the reference genome. (**C**) Genome-wide Hi-C contact map for the A4 strain (see Figure S3 for the A7 strain). Solid boxes delineate chromosome arms. Highlighted key 3D structural features (in dashed line boxes): (i) PCH of each chromosomal arm forms distinct domains; (ii) more frequent interactions occur between PCH of arms from the same chromosome than between PCH of different chromosomes; (iii) spatial clustering of PCH across chromosomes; (iv) PCH of X chromosomes exhibits clear separation from its euchromatic arm; (v) the fourth chromosome interacts with both euchromatin and PCH of other chromosomes in similar frequencies. **(D**) Number of Hi-C reads supporting 3D interactions between 5kb euchromatic windows and PCH, normalized by the uniquely mappable size of PCH for A4 strain (see Figure S4 for A7 strain). Lines/points of different colors represent interactions with PCH of a specific arm (2L, 2R, 3L, 3R, X, 4, Y) or unmapped (UNP). Insets show the total normalized contact counts for each arm. See Figures S5 and S6 for normalization using the proportion of reads, which yield consistent results.

### Reference-quality genome assemblies uncover previously uncharacterized 3D interactions involving PCH

The significantly improved PCH assemblies for the same strains used in Hi-C experiments provide an opportunity to expand our understanding of PCH 3D structure. We processed Hi-C reads using the pairtools workflow (Open2C et al. 2024) and, for this analysis, focused on Hi-C read pairs with both ends uniquely mapped to the strain-matched genome assembly (see Materials and Methods). We defined constitutive heterochromatin according to the strong H3K9me3 enrichment (Figure S2), which includes the pericentromeric regions of the second, third, and X chromosomes and the entirety of the fourth and Y chromosomes. For simplicity, we simply referred to these repressive mark-enriched regions collectively as PCH in the following analysis.

Visual inspection of the Hi-C contact maps (Figure 1C and Figure S3) confirms previous observations that PCH is hierarchically organized (Stutzman et al. 2024; Lee et al. 2020), with PCH of each chromosome arm forming distinct 3D territories (e.g., Figure 1C-i), followed by frequent interactions of PCH between arms of the same chromosome (e.g., Figure 1C-ii) and then across chromosomes (e.g., Figure 1C-iii). Additionally, interactions between PCH regions of different chromosomes (e.g., Figure 1C-iii) are significantly more frequent than contacts between PCH and euchromatic regions, supporting the established model that PCH forms a spatially segregated nuclear compartment (Janssen et al. 2018). Beyond known spatial properties of PCH, these Hi-C maps reveal a clear separation between PCH and euchromatin of the X chromosome (Figure 1C-iv), which was not detected before (Stutzman et al. 2024; Lee et al. 2020) and likely facilitated by the assembly of ∼4 Mb of additional X PCH sequence compared to previous reference genomes (Shukla et al. 2025).

We also examined the general characteristics of 3D interactions between euchromatic loci and PCH. For a given euchromatic region, a substantial fraction of its total 3D interactions with PCH occurs within the same chromosomal arm, which decays with increased linear distance between the euchromatic region and PCH (Figure 1D; Figures S4-S6). This observation is consistent with the well-established distance-dependent decay characteristic of Hi-C data, in which loci proximal in linear genomic distance exhibit higher spatial interaction frequencies than loci separated by larger linear distances (Dekker et al. 2002; Lieberman-Aiden et al. 2009). Importantly, such an observation also suggests that, while compartmentalization reduces contact probabilities between euchromatin and PCH regions, the fundamental principle of distance-dependent decay continues to shape interaction patterns across genomes. According to this distance-dependent principle, the next most frequent 3D interactions of a euchromatic region with PCH should occur with PCH from the opposite arm of the same chromosome. Surprisingly, spatial interactions between a euchromatic region and the fourth chromosome (yellow in Figure 1D) are as frequent (for the second chromosome) or more frequent (for the third chromosome) than interactions with PCH from the opposite arm (Figure 1D; Figure S4-S6). This unexpected finding may reflect the atypical epigenetic states of the fourth chromosome, which exhibits features of both euchromatin and heterochromatin (Haynes et al. 2007; Riddle et al. 2012; Riddle and Elgin 2018). Consistent with this, the fourth chromosome interacts substantially with both euchromatic and heterochromatic compartments genome-wide (Figure 1C-v). Furthermore, previously constructed 3D models positioned the fourth chromosome closer to the PCH of the third chromosome than to the PCH of the second chromosome (Lee et al. 2020), likely explaining the higher interaction frequency between the euchromatin of the third chromosome and the fourth chromosome than that between the second and fourth chromosomes. Overall, the improved PCH assemblies allowed us to confirm several previously observed PCH spatial properties while revealing new organizational patterns.

### Proposed Hi-C analysis pipeline allows allelic comparisons of 3D interactions with PCH

To identify spatial interactions between euchromatic TEs and PCH, our analysis relies on Hi-C read pairs whose two ends map to these two genomic regions. However, restricting analysis to uniquely mapped reads, a typical Hi-C analysis approach, would yield limited numbers of read pairs due to the multi-mapping propensity of reads originating from PCH, where the mappability is generally low (Figure S7). Accordingly, we developed an approach that “rescues” multi-mapped PCH reads that contain useful information for inferring euchromatin-PCH spatial proximity.

Because our goal is to estimate the total frequency of spatial interactions between euchromatic TEs and PCH, rather than identifying specific PCH regions that interact with TEs, we only need to confidently assign multi-mapped reads to PCH without resolving precise alignment coordinates. Our rescue pipeline employs a simple logic: if all possible alignment locations for a multi-mapped read fall within epigenetically defined PCH, that read is classified as PCH-derived. Conversely, if any potential alignment location maps to euchromatin, the read pair is excluded (Figure 2A; see Materials and Methods). To assess the validity and effectiveness of our proposed strategy, we first examined the genomic distribution of Hi-C read pairs whose one end mapped uniquely while the other end aligned to multiple genome locations, but all are within PCH (referred to as “rescued unique-multi read pairs” hereafter; see Table S1 for the numbers of different types of read pairs). Because of the distance-dependent decay in 3D interactions and the compartmentalization of PCH, euchromatic windows at the EU-PCH boundary are expected to exhibit the most frequent spatial interactions with PCH compared to EU regions further away from the centromere. Moreover, the majority of the interactions within heterochromatin should be confined to PCH. Consistent with these expectations, the density of rescued unique-multi read pairs significantly increases as a euchromatic window approaches PCH (Figure 2 B-i) and nearly matches that of all unique-multi read pairs within PCH (Figure 2 B-ii). These observations demonstrate that our approach successfully rescued the majority of reads that inform 3D interactions involving PCH.

**Figure 2.**
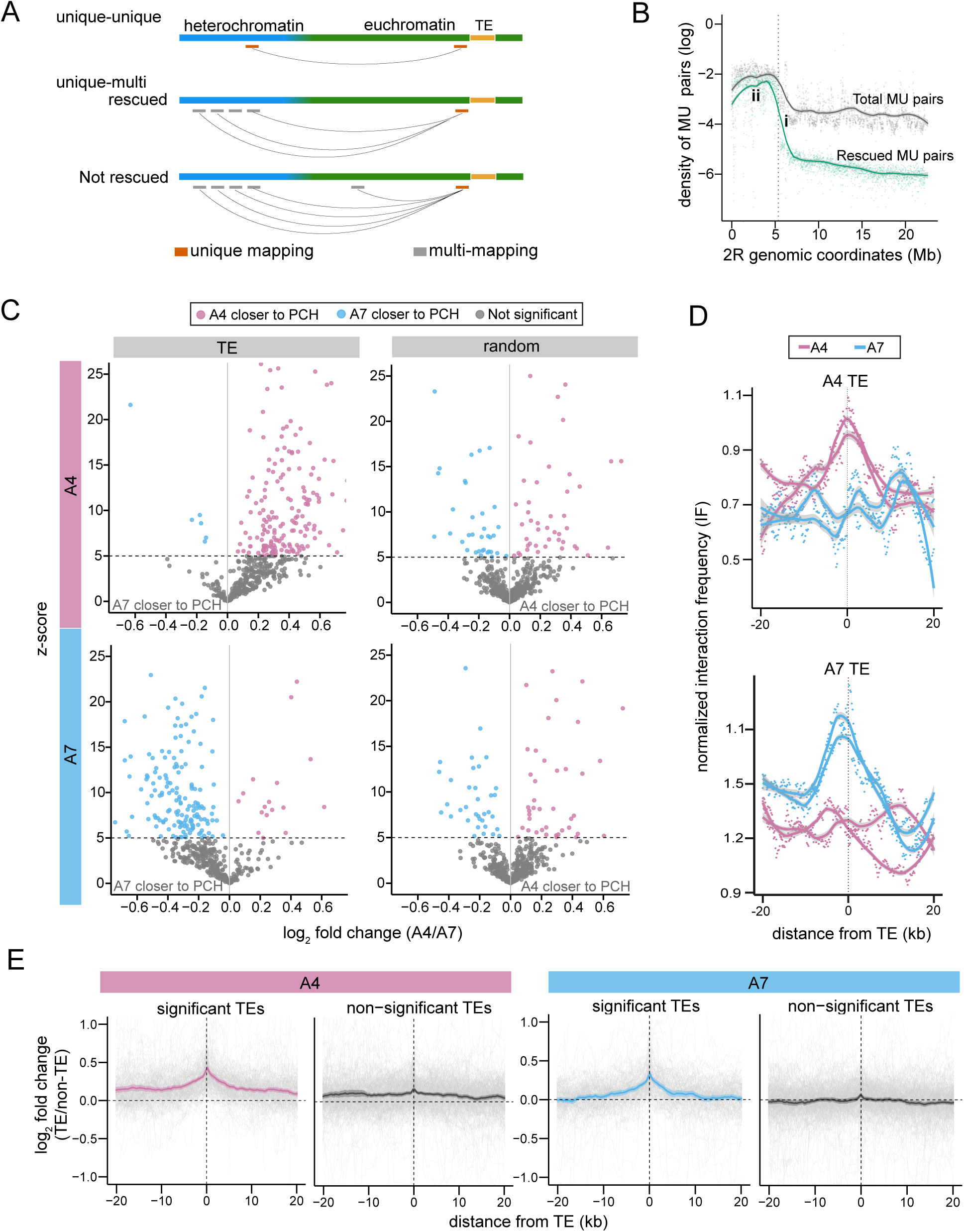
Polymorphic TEs drive varying local 3D genome organization. **(A)** Schematic illustration for the strategy of “rescuing” PCH-originating repetitive reads. Standard Hi-C analysis uses only unique-unique read pairs (top). Our pipeline rescues unique-PCH multi-mapped read pairs, where the multi-mapped ends are aligned to multiple locations that are all within PCH (middle). Pairs where the multi-mapped ends do not align exclusively to PCH are discarded (bottom). **(B)** The density of unique-multi (MU) read pairs where the unique ends aligned to a particular window of the 2R chromosome. Two lines represent total MU read pairs, irrespective of the origins of the multi-mapped end (gray) and rescued MU read pairs, where the multi-mapped end aligned exclusively in PCH (green). **(C)** Volcano plots comparing differential PCH interaction between homologous alleles for TEs (left) and randomly selected TE-free windows (right) for TE/random windows in A4 (top) and A7 (bottom). The x-axis displays the log2 fold change (log2FC) in IF values between A4 and A7 alleles, and the y-axis shows the z-scores (see text). Pink dots denote loci where the A4 alleles are significantly closer in 3D to PCH than the homologous alleles, while the blue dots represent those where the A7 alleles are closer to PCH. Gray dots denote loci without significant differences between the two alleles. TE-containing alleles exhibit a significant bias toward increased PCH interactions when compared to homologous non-TE alleles. In contrast, random control windows display a balanced distribution. **(D)** Representative examples of TE-mediated varying 3D proximity to PCH. IF values were estimated for overlapping 5 kb windows flanking TEs in A4 (top; 2L:1,926,012– 1,934,518; *roo*) and A7 (bottom; 2L:13,885,323–13,895,580; *F-element* & *Rt1b*). All of these TEs are absent in the other studied strain. **(E)** The log2 fold change of IF values between homologous alleles with and without TEs, averaged separately for TEs that exhibit significant 3D interactions with PCH versus those that do not.

We used Hi-C read pairs with both ends mapped uniquely, as well as rescued unique-multi read pairs, to identify TEs that significantly enhance 3D interactions with PCH by comparing homologous alleles with and without TEs. The TE-free allele served as a null control for the expected PCH interaction frequency, which arises primarily from interactions of a focal euchromatic region with the PCH of the same chromosome (including both same- and opposite-arm PCH) as a consequence of distance-dependent decay. To enable quantitative comparisons between with- and without-TE alleles, we developed a two-stage normalization strategy across replicates and genotypes (see Materials and Methods for details). Briefly, within a replicate, the first round normalizes the number of Hi-C read pairs supporting euchromatin-PCH 3D interactions by the total number of read pairs with at least one end mapped to the euchromatic window, an approach that accounts for differences in sequencing depth and sequence mappability across genomic regions. The second round addresses differences in Hi-C library complexity between samples and replicates by using PCH interaction frequencies of local genomic regions (500 kb surrounding each focal window) to derive sample-specific scaling factors. These approaches normalized interaction frequencies toward a common baseline and yielded a normalized “Interaction Frequency” (IF), with higher IF values indicating more frequent 3D interactions between the focal euchromatic region and PCH. For randomly selected euchromatic windows lacking TEs in either strain, we found no systematic bias in IF values between homologous alleles, confirming that our normalization strategy successfully accounts for technical variation between samples and replicates (Figure S8; see Materials and Methods for details).

### Polymorphic TEs drive varying local 3D genome structures

To test whether polymorphic TEs drive varying local 3D structures through spatial interactions with PCH, we compared homologous sequences with and without TEs using the normalization framework outlined above. Comparisons between near-identical homologous alleles allow excluding the confounding influence of local sequence context that could not be controlled in previous analyses using randomly sampled genomic regions within the same strain as controls (Lee et al. 2020). Because our goal is to assay each euchromatic TE separately, we need to distinguish Hi-C read pairs originating from individual TE insertions. Using Hi-C reads aligning directly to TEs is not feasible due to their repetitive nature. Instead, we rely on the fact that TEs and neighboring sequences are immediately adjacent on the linear chromosome, and spatial interactions between TEs and PCH should bring the neighboring sequences into proximity of the PCH as well. We therefore analyzed Hi-C read pairs whose one end aligned uniquely to 5kb windows flanking the left and right boundaries of a TE insertion, or to a 10kb homologous window in the strain lacking that TE insertion.

We annotated TEs in the reference-quality assemblies and performed structural variant calling between genomes to define homologous windows in the reciprocal genome (see Materials and Methods). This yielded 498 and 471 TEs that are unique in A4 and A7, respectively. We then quantified IF values for homologous windows with and without a particular TE and used these values to calculate z-scores (differences in mean IF values between homologous alleles, normalized by their standard deviation; see Materials and Methods). We performed analyses using either (1) only uniquely mapped Hi-C read pairs or (2) uniquely mapped plus rescued multi-mapped Hi-C read pairs. The latter approach recovers significantly more Hi-C read pairs per TE (median of 158 vs. 65; *Mann-Whitney U test, p* < 10⁻¹⁶) and results in smaller variance of IF estimates between replicates (coefficient of variation 0.040 vs. 0.057; *Mann-Whitney U test, p* < 10⁻¹⁶). Because the two approaches yield consistent results and the latter approach has higher statistical power, we present analyses combining uniquely mapped and rescued multi-mapped reads in the main text and those based only on uniquely mapped reads in the supplementary figures (Figures S9 and S10).

We found that the presence of TEs leads to larger IF values when compared to the homologous alleles without TEs (Figure 2C, left column; Figure S9). With a significance threshold of z-score > 5 or *p-value* < 0.05, 37.12% of TE-containing alleles (39.38% in A4 and 34.75% in A7) are significantly closer to PCH than the homologous TE-free alleles. This is in stark contrast when performing the same analysis using randomly sampled TE-free windows with matching chromosomal distributions (Figure 2C, right column; Figure S9), with only 9.12% (9.83% in A4 and 8.37% in A7) of the windows being significant (*Fisher’s Exact test*, *p <* 10^-16^ for all comparisons). With a more stringent threshold (z-score > 5 and log2 fold change > 0.2, corresponding to IF values at least 1.15-fold higher in TE-containing versus TE-free alleles), we found even more dramatic differences: 32.15% (A4) and 29.00% (A7) of TEs versus only 5.32% (A4) and 5.36% (A7) of random windows were significant. In two representative examples, both replicates of the TE-containing alleles show a marked increase in IF values as windows approach the TE insertion sites, while IF values of the TE-absent allele remain low and nearly flat across windows (Figure 2D). Notably, for these example TEs, TE-mediated spatial proximity to PCH extends well beyond the 5 kb flanking windows we used to identify TEs that drive significant polymorphic 3D structures. We thus investigated the extent, along a linear chromosome, of TE-mediated PCH proximity by averaging IF values around all analyzed euchromatic TEs, revealing that PCH proximity typically extends ∼10 kb from TE insertion sites (Figure 2E; Figure S10). Overall, our findings provide the first strong genome-wide evidence that polymorphic TEs drive varying local 3D genome structures.

### TEs engaged in 3D interactions with PCH differ from other TEs in chromosome locations and biological properties

Not all analyzed euchromatic TEs drive 3D proximity to PCH when compared to homologous TE-free sequences, and identifying the biological attributes of these TEs could help elucidate the potential functional and evolutionary consequences for both host genomes and TEs. We investigated the genomic locations of TEs showing significant 3D interactions with PCH (z-score > 5), examining chromosome arms, linear positions within arms, and local chromatin environment. While the proportion of significant TEs on each autosomal arm does not significantly differ (Figure 3A), TEs on the X chromosome are less likely to show significant spatial proximity to PCH in one of the two strains (A4: 40% (autosome) vs 39% (X) of TEs, *Fisher’s Exact Test, p* = 0.91; A7: 37% (autosome) vs 26% (X) of TEs, *Fisher’s Exact Test, p* = 0.06). This observation could result from the more diffuse associations between X euchromatic regions and autosomal PCH (Figure 2D). Alternatively, we may simply have lower statistical power to detect X-linked TE-PCH interactions due to reduced X chromosome sequencing coverage in our mixed-sex embryo samples.

**Figure 3.**
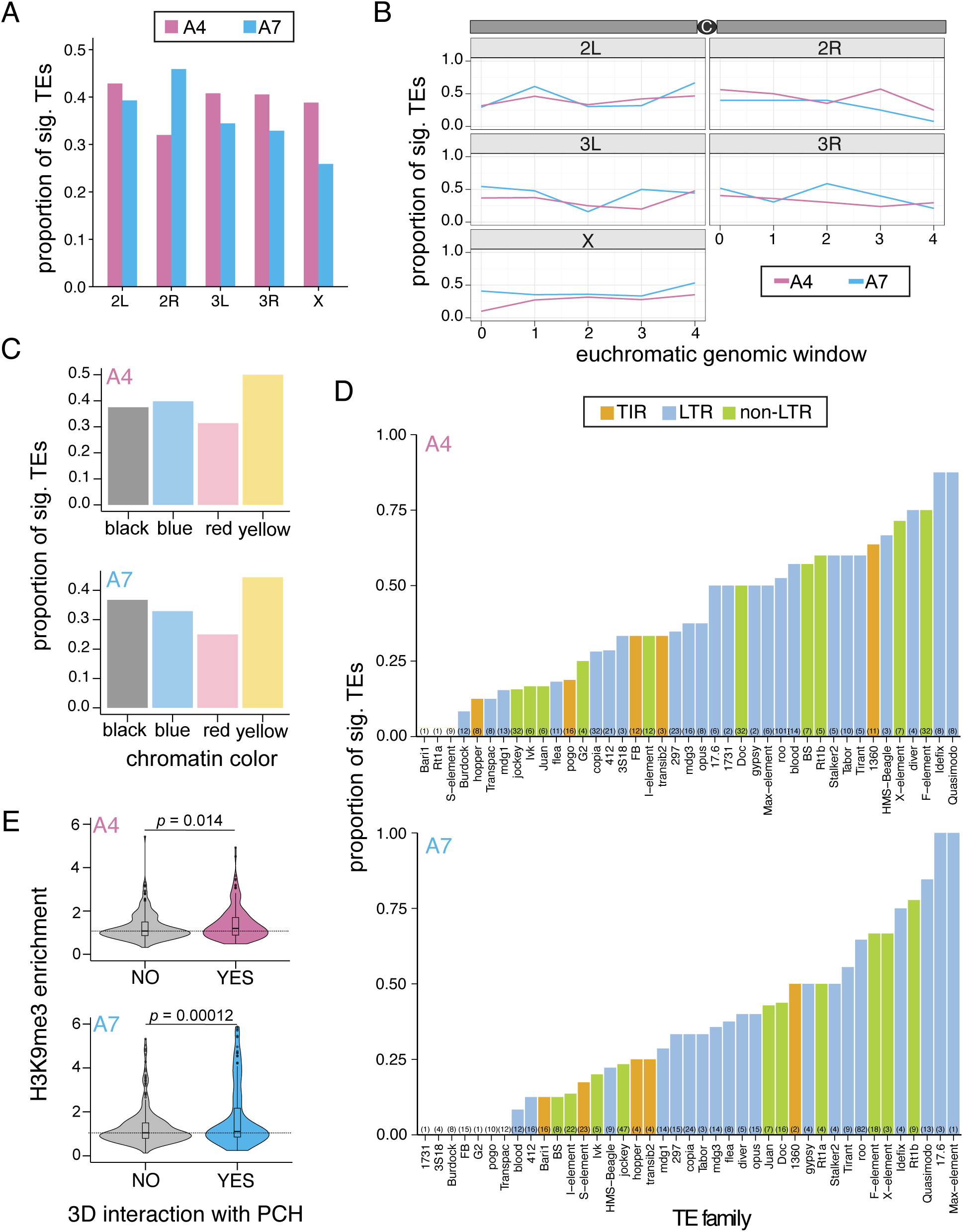
Genomic distribution and biological properties of TEs engaged in PCH 3D interactions. **(A and B)** Proportion of TEs with significant PCH 3D interactions across chromosome arms (A) and genomic windows (∼4-5 Mb) along each chromosome arm (B) for A4 (pink) and A7 (blue). **(C)** Proportion of TEs with significant PCH 3D interactions for genomic regions of varying chromatin state. **(D)** Proportion of TEs with significant PCH 3D interactions for each studied TE family, colored by class (DNA, LINE, LTR). **(E)** Violin plots comparing H3K9me3 enrichment in the 5 kb flanking regions of TEs with and without significant PCH interactions.

When TEs are grouped by their linear position along each chromosomal arm (∼4–5 Mb bins), the proportion showing significantly enhanced proximity to PCH compared to their TE-free homologs remains similar across all bins (Figure 3B), suggesting that TE-mediated PCH interactions are largely independent of linear genomic distance from PCH. This observation contrasts with previous findings that TEs closer to PCH on a linear chromosome are also more likely to engage in TE-PCH 3D interactions (Lee et al. 2020). Such a discrepancy likely arose because the previous investigation was based on a single Hi-C experiment and used randomly sampled genomic regions to generate “null distributions” for identifying TEs showing significant 3D interactions with PCH, an approach that may fail to control for the distance-dependency of spatial interactions and unrecognized confounding factors in local sequence context (Lee et al. 2020). It is worth noting that the absolute interaction frequency with PCH (without comparison to TE-free homologs) still increases as TEs approach PCH boundaries due to distance-dependent decay. Our comparisons between homologous alleles with and without TEs control for this baseline positional effect, which could not be controlled for in previous studies(Lee et al. 2020).

We investigated whether TEs in genomic regions with different chromatin states exhibit varying propensities for significant 3D interactions with PCH. We utilized previously defined chromatin states generated from enrichment profiles of multiple histone modifications and chromatin-interacting proteins in an embryonic cell line (Filion et al. 2010). This classification segments the genome into five types: two active states (yellow for housekeeping genes and red for developmentally regulated genes), two classic repressive states (green for H3K9me2/3-enriched constitutive heterochromatin and blue for Polycomb-mediated facultative heterochromatin), and black chromatin, a novel repressive state lacking classic heterochromatin marks. Although these chromatin data were collected from a different genotype lacking the TEs we studied, we used them to define broadly conserved chromatin environments and test whether TEs in certain chromatin contexts show different propensities to interact with PCH. TEs enhancing 3D PCH interactions are not significantly overrepresented in repressive blue or black chromatin states (Figure 3C; insufficient observations for green chromatin). Intriguingly, TEs in yellow chromatin, an active state enriched with H3K36me3, show a trend toward more frequent significant 3D interactions (Figure 3C; *Fisher’s Exact Test,* odds ratio = 1.68, *p* = 0.19 (A4) and odds ratio = 1.52, *p* = 0.30 (A7)), though these differences did not reach statistical significance.

We then investigated biological attributes of TEs that may have driven their propensity to spatially interact with PCH. Among 42 TE families analyzed, 100% of them have at least one TEs showing significant 3D interactions with PCH in either strain, and these TE families are from all major super families (Figure 3D). Nevertheless, TEs that transpose through an RNA intermediate (Class I) are more likely to significantly interact with PCH than those move through a DNA intermediate (Class II; proportion of significant TEs in A4: 41% (RNA) vs 24% (DNA), *Fisher’s Exact test, p* = 0.18; in A7: 39% (RNA) vs 13% (DNA), *Fisher’s Exact test, p* = 1.8×10^-5^). Breaking down TEs into sub-types still identified both RNA-based types of TEs, LTR and non-LTR, are significantly more likely to spatially interact with PCH than DNA-based TIR TEs (Figure S11).

Because it was hypothesized that TE-PCH 3D interactions are mediated by H3K9me2/3 and recruited HP1a (Lee et al. 2020), we used our previously published H3K9me3 data from the same embryonic stage and genotype (Huang et al. 2025) to compare H3K9me3 enrichment between TEs with and without significant PCH 3D interactions. However, the repetitive nature of TEs and the reliance of standard histone modification profiling on short-read sequencing (Kaya-Okur et al. 2019) prevented us from estimating H3K9me3 enrichment within TEs. Instead, we leveraged the widely documented phenomenon that repressive marks at silenced euchromatic TEs spread to nearby sequences (Choi and Lee 2020) and estimated H3K9me3 enrichment in the 5 kb windows flanking each TE (left and right; the same windows used for Hi-C analysis). TEs interacting with PCH have significantly higher enrichment of H3K9me3 (Figure 3E: A4: median enrichment level of H3K9me2, 1.19 vs 1.07, *Mann-Whitney U test, p =* 0.014; A7: median enrichment level of H3K9me2, 1.25 vs 1.07, *Mann-Whitney U test, p =* 1.2×10^-4^). Because longer TEs, when epigenetically silenced, represent larger stretches of heterochromatin, we predict they may more readily facilitate TE-PCH 3D interactions. Consistently, TEs interacting with PCH are significantly longer than other TEs (Figure S12). It is worth noting that longer TEs also tend to result in higher enrichment of H3K9me3 in flanking sequences (Lee 2015; Huang et al. 2022), an observation we also found (*Spearman rank correlation coefficient* A4: *ρ* = 0.26, p = 4.9×10^-9^; A7: *ρ* = 0.42, p < 10^-16^). Nevertheless, logistic regression analysis that includes both TE-mediated enrichment of H3K9me3 and TE length found that both indices contribute to the propensity of TEs involved in 3D interactions with PCH (3D interaction or not ∼ H3K9me3 enrichment + length; A4: H3K9me3 coefficient = 0.22, *p* = 0.13; length coefficient = 9.5×10^-5^, *p* = 0.00041. A7: H3K9me3 coefficient = 0.13, *p* = 0.0023; length coefficient = 10^-4^, *p* = 0.0027). Also, although RNA-based TEs are commonly observed to lead to higher enrichment of H3K9me3 (Lee 2015; Huang et al. 2022; Lee and Karpen 2017), regression analysis within class or type of TEs consistently found that TEs with PCH 3D interactions have higher enrichment of H3K9me3 (Table S2). Overall, our analyses indicate that TEs significantly engaged in PCH 3D interactions differ from other TEs in their genomic locations (autosomal) and biological attributes (RNA-mediated transposition, longer length, and H3K9me3 enriched).

### 3D interactions between TEs and PCH have functional and evolutionary consequences

PCH is highly enriched with chromatin proteins that are associated with the suppression of various genome functions, including transcription as well as recombination and repair (Janssen et al. 2018). For the latter, due to the repetitive sequence composition of PCH, DNA breaks within the heterochromatin domain are dynamically moved outside that domain for repair, leading to suppressed aberrant recombination within the PCH domain (Chiolo et al. 2011; Janssen et al. 2016). We postulated that TE-mediated proximity to PCH could have similar impacts on the distribution of recombination events. We leveraged our previously generated maps of crossovers, one of the two major recombination products, in one of the two strains used in this study (Huang et al. 2025) and found marginal statistical support for reduced crossover occurrence upon TE-mediated proximity to PCH (A7: mean number of crossovers: 0.02 (with 3D interactions) vs. 0.08 (without 3D interactions); *Mann-Whitney U test, p* = 0.09).

In addition, spatial proximity to the PCH was found to suppress the expression of euchromatic genes mispositioned into the PCH domain (Harmon and Sedat 2005; Dernburg et al. 1996; Csink and Henikoff 1996). We predicted that spatial interactions between polymorphic TEs and PCH could similarly bring local euchromatic regions into PCH proximity, leading to reduced expression of TE-adjacent alleles when compared to homologous TE-free alleles. To test this prediction, we generated transcriptome data from the same embryonic stage and strain (see Materials and Methods). To compare expression between homologous alleles with and without adjacent TEs, we ranked gene expression from highest to lowest within each strain and used these ranks to calculate z-scores between with- and without-TE alleles (mean difference between strains normalized by standard deviation). A positive z-score indicates lower expression of TE-containing alleles relative to their TE-free counterparts. For genes within 5 kb of TEs that lack 3D interactions with PCH but are enriched for H3K9me3, we found no significant difference in z-scores compared to other genes, indicating no TE-mediated changes in expression (Figure 4A, A4: median z-score with and without H3K9me3, 0.34 vs 1.47, *Mann-Whitney U test, p =* 0.21; A7: median z-score with and without H3K9me3, -0.15 vs 0.24, *Mann-Whitney U test, p =* 0.17). This observation is consistent with previous observations that TE-mediated local enrichment of repressive marks alone does not have a directional influence on the expression of adjacent genes ((Lee and Karpen 2017; Huang et al. 2022) and reviewed in (Choi and Lee 2020; Kelleher et al. 2020)). Interestingly, genes whose adjacent TEs show 3D interactions with PCH and are enriched for H3K9me3 tend to have higher z-scores, with the difference being significant for one of the two strains (Figure 4A; A4: median z-score 0.60 (with 3D interactions) vs. 0.34 (without 3D interactions), *Mann-Whitney U test, p =* 0.28; A7: median z-score 0.49 (with 3D interactions) vs. -0.15 (without 3D interactions), *Mann-Whitney U test, p =* 0.025). This observation indicates that TEs with PCH 3D interactions lower the expression of adjacent genic alleles when compared to homologous TE-free alleles. Notably, we observed a case in which three clustered TEs are associated with suppressed gene expression over a 60 kb genomic region encompassing seven genes, likely mediated through TEs’ proximity to PCH (Figure 4B).

**Figure 4.**
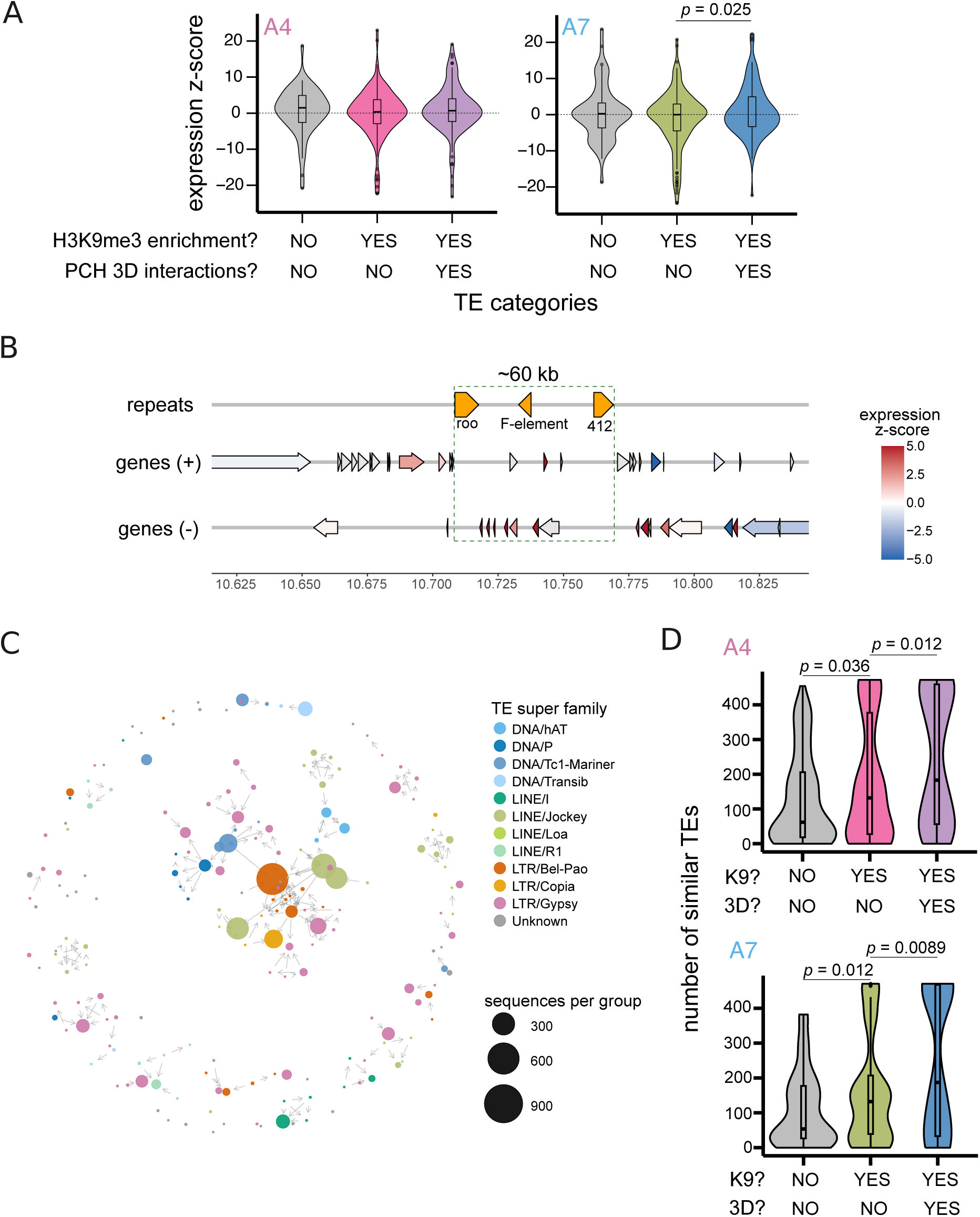
Functional and evolutionary consequences of TE-mediated 3D interactions with PCH. **(A)** z-scores comparing the expression rank of homologous genic alleles with and without flanking TEs. A positive z-score indicates that the with-TE allele has lower expression than the homologous TE-free allele. TEs are classified based on H3K9me3 enrichment and, for those with enrichment, whether they exhibit significant PCH 3D interactions. **(B)** Representative example for the suppressive effects of TE-PCH 3D interactions on nearby gene expression. A ∼60 kb genomic region on chromosome 2L harbors a cluster of three TEs (orange triangles): *roo* (z-score for PCH 3D interaction = 3.16), *F-element* (z-score = 36.45), and *412* (z-score = 0.82). Genes are depicted as arrows to indicate transcriptional direction and are colored based on expression rank differences between homologous alleles. Red shading denotes lower expression in the TE-containing allele, while blue denotes higher expression. Notably, genes within this ∼60kb region show predominantly reduced expression in the TE-containing allele, contrasting with the variable up- and down-regulation of genes immediately outside this region. **(C)** Connection graph between TE insertions based on sequence similarity, colored by TE superfamilies. Nodes represent clusters of TEs with similar sequences, with node size proportional to cluster size. Most nodes (99.3%) are assigned to specific TE superfamilies. Directed edges denote one-way sequence similarity, indicating nesting of one TE (arrow origin) within another TE (arrow end). **(D)** Number of similar TEs for three categories of TEs: (1) without enrichment of H3K9me3 nor PCH 3D interactions, (2) H3K9me3-enriched TEs without PCH 3D interactions, and (3) H3K9me3-enriched TEs with PCH 3D interactions.

Under the general expectation that gene expression levels are at fitness optima and genetic variants that alter gene expression are under purifying selection (Bedford and Hartl 2009), we predicted that reduced gene expression mediated by TE-PCH 3D interactions should impair individual fitness. Assuming a constant transposition rate, TEs that are more strongly selected against should have low population frequencies and/or be young, a commonly adopted approach for inferring selection strength against TEs (reviewed in (B Charlesworth and Langley 1989; Lee and Langley 2010; Barrón et al. 2014). We estimated the population frequencies of TEs in an African population of *D. melanogaster* (Lack et al. 2015), which represents the likely ancestral range of the species and is less likely to have experienced recent demographic changes that could skew the distribution of allele frequencies. While TEs with 3D interactions have lower population frequencies than other TEs (Figure S13), the generally low frequencies of studied TEs lead to a lack of statistical power. To infer relative TE age, we leveraged the fact that each TE copy begins accumulating mutations upon insertion into a new genomic location, such that younger TEs have accumulated fewer mutations and thus remain more similar to other copies. We leveraged the highly contiguous assemblies of strains sampled across the global genetic diversity of the species (Chakraborty et al. 2019) and used Pannagram (Igolkina et al. 2025a) to construct “connection” among TE insertions based on sequence similarity (Figure 4C (color coded by superfamily) and Figure S14 (color coded by TE family); see Materials and Methods). Consistent with our prediction, TEs with PCH interactions tend to have significantly more similar TEs (85% similarity) than those without such interactions (Figure 4D; A4: median 183 vs. 132, *Mann-Whitney U test, p* = 0.012; A7, median 187 vs. 132, *Mann-Whitney U test, p =* 0.0089). This observation indicates that TEs with PCH 3D interactions tend to be younger, which indicates they may have experienced stronger purifying selection. To exclude the possibility that our observed differences in TE age are driven by variation in recent transposition activity between types of TEs (Blumenstiel et al. 2014; Bergman and Bensasson 2007), rather than differential selection strength against TEs, we conducted the same analysis within TE types, yielding qualitatively consistent results (Figure S15). Overall, we found that polymorphic 3D structures driven by TE-PCH spatial interactions may reduce gene expression of TE-adjacent alleles relative to TE-free homologous alleles, and this functional impact could impose stronger selection against TEs engaged in 3D interactions with PCH.

## Discussion

Understanding the molecular drivers of variation in 3D genome organization is critical for explaining how functionally consequential differences in 3D genome architecture arise and evolve. Structural variants, including deletions and duplications, have been identified as genetic determinants that contribute to variation in 3D genome structure (Gilbertson et al. 2024) or to the disruption of such structure (Spielmann et al. 2018; Shanta et al. 2020). These variants generate varying 3D genome organizations primarily by altering the location, abundance, or orientation of boundary elements (Spielmann et al. 2018), which are DNA sequences that establish chromatin loops and higher-order topologically associated domains (TADs), the fundamental organizational units of spatial genome structure (Dekker and Mirny 2024). TEs, which are also a type of structural variant under the broad definition, were similarly implicated in mediating 3D genome organization, mainly based on observations that specific TE families are enriched at TAD boundaries (Bourque et al. 2018; Schmidt et al. 2012). Subsequent studies corroborated these associations and, through direct experimental manipulation, demonstrated that TEs can function as boundary elements that shape 3D genome organization (Lawson et al. 2023; Fueyo et al. 2022). Given the substantial between-species divergence in TE profiles, both in composition and abundance (Elliott and Gregory 2015; Osmanski et al. 2023), this TE-mediated 3D organization is expected to contribute to interspecific differences in genome spatial structure. Indeed, TEs were found to account for an appreciable proportion of variable chromatin loops and TAD boundaries across mammals (Choudhary et al. 2023; Diehl et al. 2020; Choudhary et al. 2020).

However, these previous studies on TEs’ role in 3D genome organization have been limited to interspecies comparisons. Whether the substantial variability of TEs within species (e.g., (Cridland et al. 2013; Rech et al. 2022)) generates polymorphic 3D genome architecture remains largely unexplored. Here, we demonstrated, for the first time to our knowledge, that polymorphic TEs drive variation in local 3D genome structure within species. We achieved this by overcoming several technical challenges, including generating reference-quality genome assemblies that enable assaying genetic variation within repetitive PCH of studied strains and developing a new Hi-C analysis framework that augments identification of PCH reads while enabling between-strain comparisons. We found that nearly 40% of the polymorphic TEs enhance spatial interactions with PCH when compared to TE-free homologs, producing between-strain variation in local 3D structures. Given previous findings that more than 3,000 TEs segregate at low population frequencies in a sample of five genomes (Rech et al. 2022), and considering the observed extent of these TE-mediated effects (∼10 kb on each side of a TE; Figure 2E), our observations suggest that 20% of the 120 Mb euchromatic genome would exhibit TE-mediated variation in 3D proximity to PCH even in this limited sample size. Interestingly, our allele-specific comparisons, which control for baseline distance-dependent decay in spatial interactions, reveal that the propensity for TEs to spatially interact with PCH remains consistent across chromosome arms, regardless of the linear distance between TEs and PCH. Moreover, while TEs exhibiting significant 3D interactions with PCH are more likely to be RNA-based and longer in length, their most defining feature is the enrichment of repressive H3K9me3. These observations suggest a chromatin-based mechanism that stands in sharp contrast to prior mechanisms in which TEs shape 3D genome structures through specific DNA sequence characteristics of particular TE families that act *in cis* (e.g., (Thybert et al. 2018; Choudhary et al. 2023), reviewed in (Lawson et al. 2023)). The observed 3D interactions between polymorphic TEs and PCH extend broadly to insertions from diverse TE families across genome locations (Figure 3B and 3D), establishing polymorphic TEs as a potent and general driver of variation in genome organization between individuals.

The identified TE-mediated polymorphism in 3D proximity to PCH likely has functional consequences, as suggested by our observation that PCH-interacting TEs tend to reduce the expression of adjacent genic alleles compared to TE-free homologs. This observation echoes previously documented phenomena where spatial mislocalization of euchromatic genes into heterochromatin domains leads to gene silencing (Harmon and Sedat 2005; Dernburg et al. 1996; Csink and Henikoff 1996). It is worth noting that both the Hi-C and RNA-seq data were generated from bulk embryos. The suppressive impacts of TE-mediated 3D proximity to PCH on nearby gene expression could be more substantial in individual cells where this spatial repositioning actually occurs than our population-averaged measurements suggest. Interestingly, among TEs that induce H3K9me3 enrichment at neighboring genes, only those that also exhibit significant 3D interactions with PCH result in reduced expression of adjacent genes (Figure 4A). Multiple studies across taxa have similarly found that TE-induced enrichment of repressive marks does not predominantly reduce neighboring gene expression (reviewed in (Choi and Lee 2020; Kelleher et al. 2020), but see (Coronado-Zamora and González 2025)). These observations indicate that the enrichment of repressive epigenetic modifications alone is insufficient to suppress neighboring gene expression and that spatial interaction with PCH is additionally required. Such a requirement for both chromatin modification and spatial repositioning may be analogous to the action of enhancers, where the deposition of specific histone modifications alone cannot affect gene expression without spatial interaction with promoters (e.g., (Kragesteen et al. 2018; Chen et al. 2024; Golov et al. 2024)). Importantly, our observation that TEs show similar propensities to engage in spatial interactions with PCH regardless of their linear distance from PCH suggests that these interactions may transcend chromosomal substructures (e.g., TADs) that typically constrain gene regulatory interactions. This raises the possibility that TE-mediated variation in gene expression through 3D interactions with PCH may extend well beyond the immediate TE insertion site.

TEs involved in 3D interactions with PCH tend to be evolutionarily young, signatures of being subject to strong purifying selection, indicating that TE-mediated changes in 3D genome structures likely impose fitness costs. These costs may arise not only from the reduced neighboring gene expression discussed above, but also from impacts on other genome functions that depend on proper spatial localization, such as DNA repair and recombination (Chiolo et al. 2011; Janssen et al. 2018), although we only obtained marginal statistical support for the latter. While all identified TE-PCH 3D interactions in our study are most likely deleterious due to the very low (<2%) population frequencies of TEs involved, it remains an intriguing question whether this mechanism might occasionally confer beneficial functions, undergo positive selection, and drive the evolution of 3D genome structures between species. Cases of adaptive TEs have been documented (Feschotte 2008; Chuong et al. 2017), with most contributing to TE-derived protein-coding sequences (Cosby et al. 2021; Pasquesi et al. 2024) or regulatory elements that upregulate nearby genes (Daborn et al. 2002; Hof et al. 2016). In contrast, definitive cases of adaptive TE-mediated gene downregulation are scarce and, when documented, typically involve complex regulatory changes affecting multiple genes through both up- and down-regulation (e.g., (Guio et al. 2014; González et al. 2009)). Nevertheless, several observations suggest that TE-mediated gene silencing through spatial interactions might occasionally be beneficial. Reduced gene expression can be adaptive in certain contexts (Groen et al. 2020), and population surveys have identified high-frequency TEs (>15%) associated with reduced expression of adjacent genes (Rech et al. 2022). Also, gene silencing mediated through spatial localization to repressive heterochromatic compartments is crucial for several biological functions, such as ensuring the expression of only one receptor gene out of hundreds per olfactory neuron in mammals (Clowney et al. 2012; Magklara et al. 2011). TE-mediated 3D repositioning of neighboring genes to the repressive PCH domain would represent a conceptually similar mechanism that could achieve adaptive gene silencing.

Importantly, TE-mediated silencing via spatial repositioning can contribute to adaptive evolution only if it varies between populations and species. In addition to the substantially different insertion locations and compositions of TEs, the chromatin-based nature of TE-PCH interactions suggests another source of variation. The enrichment of repressive marks at euchromatic TEs correlates with the expression of genes involved in heterochromatin function across species (Huang et al. 2022; Lee and Karpen 2017). Varying expression levels and, potentially, protein sequences of genes that modulate heterochromatin environment (Lin et al. 2024a) could drive varying enrichment of repressive marks at euchromatic TEs and, consequently, their propensity to spatially interact with PCH. Through TE-PCH 3D interactions, variation in both TE profiles and chromatin environment may play critical, yet previously unrecognized, roles in shaping the evolution of 3D genome organization, with functional consequences extending beyond the immediate TE neighborhood.

## Materials and Methods

### Drosophila husbandry

A4 and A7 strains from DSPR were reared on standard media at 25 °C with a 12/12 dark/light cycle with controlled humidity. Mixed sex 16-18-hour embryos were collected on standard apple juice plates (for Hi-C and RNA-seq experiments). Adult male flies were collected for PacBio HiFi sequencing.

### Pacbio HiFi Sequencing and Assembly

High Molecular Weight (HMW) DNA was extracted from ∼200 adult males for the A7 strain following the protocol from (Shukla et al. 2025). HMW DNA samples were sheared via gTUBE (Covaris) and prepared into libraries using the SMRTbell Express Template Prep Kit 2.0 (Pacific Biosciences, Menlo Park, CA) with a final size selection step, following the manufacturer’s recommendation. Sequencing was conducted on a Pacific Biosciences Sequel II platform at the UC Irvine Genomics High-Throughput Facility. The assembly was constructed according to the methods detailed in (Shukla et al. 2025). Briefly, hifiasm 0.16.1 (Cheng et al. 2021) was used to assemble raw HiFi reads into contig assemblies. The contig-level assemblies were scaffolded using reference (ASM4260644v1) assisted scaffolding to arm-level scaffolds. PacBio HiFi assembly from (Shukla et al. 2025) was used for the A4 strain.

### Identification of polymorphic TEs

The *D. melanogaster* consensus TE library was obtained from the Bergman Lab GitHub repository (v10.2; https://github.com/bergmanlab/drosophila-transposons) and used as input for RepeatMasker (v4.1.2-p1) (Smit et al. 2015) to annotate TEs in the A4 and A7 assemblies. Adjacent TE annotations of the same family separated by less than 200 bp were merged. TE annotations were filtered to retain elements with a size greater than 1,000 bp. Annotated TEs within 5 kb of each other were consolidated into single loci when investigating their 3D proximity to PCH. To identify genomic breakpoint coordinates for these TEs in the reciprocal genome and evaluate whether they are absent in the other strain, we employed two structural variant calling pipelines to identify deletions between A4 and A7 genomes bidirectionally: (1) minimap2 (v2.25) (Li 2018, 2021) for whole-genome alignment, followed by SV detection with paftools.js, and (2) wfmash (v0.10.5) (Marco-Sola et al. 2021) for alignment, followed by SyRI (v1.6.3) (Goel et al. 2019) for structural variant calling. Results from both pipelines were consolidated to maximize detection coverage. TEs were excluded if their corresponding breakpoint in the reciprocal genome was within 5 kb of any annotated TE.

### Hi-C Library prep and Sequencing

Hi-C experiment was performed using Arima Genomics (Carlsbad, California) Arima-HiC 2.0 kit (Arima High Coverage) with 16-18-hour mixed sex embryos, with two biological replicates for each genotype. This protocol is optimized for uniform coverage across genomes and reduced sequence-specific biases by employing a four-enzyme cocktail of frequent cutters (specifically 4-bp and degenerate 5-bp motifs) rather than a single restriction enzyme. Proximally-ligated DNA was then fragmented using Covaris, followed by size selection using AMPure beads and Biotin enrichment, and library prep using Swift Biosciences Accel-NGS 2S Plus DNA library kit, all following Arima Genomics protocols. Produced libraries were sequenced on Illumina NovaSeq.

### Processing of Hi-C data

Processing of Hi-C reads was conducted following the guidelines of the *pairtools* software suite (v1.0.2) (Open2C et al. 2024). Briefly, paired-end Illumina reads were aligned to their respective PacBio HiFi genome assemblies using BWA-MEM (v0.7.17-r1188) (Li 2013) with the -SP and additional -Yq flags to ensure appropriate handling of supplementary alignments that are essential for the rescuing of PCH multi-mapped reads (see below). To ensure comprehensive representation of the PCH region, the genome assemblies were augmented *a priori* with canonical *D. melanogaster* satellite repeat sequences to account for potential incomplete satellite representation in the assemblies. Because the Hi-C experiment was conducted using multiple restriction enzymes, an appreciable fraction of read pairs contained more than one ligation junction, leading to split or chimeric alignments that are discarded by typical Hi-C analysis. To maximize data retention and to rescue Hi-C read pairs with more than one ligation event, aligned reads were processed and classified using *pairtools parse* with the *--walks-policy all* parameter. We retained Hi-C read pairs classified as “UU,” indicating pairs where both reads mapped uniquely (MAPQ>0), and “MU,” indicating pairs with one uniquely mapped read (MAPQ>0) and one multi-mapping read (MAPQ=0), for the subsequent analyses. A summary of the number of read pairs at each data processing stage is in Table S1. PCR duplicates were removed using *pairtools dedup*. HiCExplorer (Ramírez et al. 2018) was used to perform quality control and assess reproducibility between biological replicates, confirming that the replicates exhibited high concordance (Figure S16).

### Rescuing of multiple-mapped PCH reads

Multi-mapping (M) reads were defined by a MAPQ score of 0, indicating equally optimal alignments at multiple genomic locations. MU read pairs were isolated from the *pairtools* output file and deduplicated using a custom Python script due to the fact that *pairtools dedup* only deduplicates UU read pairs. To identify multi-mapped PCH reads, we used *bwa mem* with the *-aq* options to output all alignments and only retain reads where all mapped locations are within the epigenetically defined PCH region (see below). A final filtering step was applied to the set of rescued EU-PCH pairs, requiring the unique euchromatic end to have a MAPQ score > 30 in order to minimize potential false positives.

### Estimation of the Interaction Frequency (IF) index

To enable quantitative comparison of EU-PCH interactions across biological replicates and strains, we developed a two-stage normalization strategy accounting for sequencing depth and library complexity. In the first stage, we normalized for local sequencing depth variation. For each 5 kb euchromatic window, we used the total count of Hi-C read pairs whose both ends uniquely mapped (UU pairs) and at least one end mapped to that window as a proxy for local sequencing depth. The count of EU-PCH read pairs in each window was then divided by its local UU pair count to generate an initial normalized interaction frequency.

During the initial quality control, we noticed that A7 samples captured higher proportions of long-range interactions than A4 samples (Table S1), leading to a higher proportion of reads supporting PCH 3D interactions for the A7 allele than the A4 allele, even for randomly selected TE-free windows (Figure S8). Aligning Hi-C data to the iso-1 reference genome assembly, which mitigates potential artifacts arising from differences in PCH representation between our strain-specific assemblies, found similar biases (Figure S17). To account for differences in library complexity, we implemented a second, localized normalization step. First, using depth-corrected values from stage one, we calculated normalized interaction frequencies (IF) for non-overlapping 5 kb euchromatic windows in each replicate, excluding windows overlapping annotated TEs. We then applied localized normalization: for each focal window, we collected IF values from the surrounding 500 kb region (50 windows upstream and 50 windows downstream). We determined the median IF value within this local window set for each sample independently and identified the minimum median IF value across all samples. A sample-specific scaling factor was derived by dividing this minimum median IF value by the sample’s median IF, and this factor was applied to adjust the focal window’s IF. For windows adjacent to TE insertions, the four immediately flanking windows (20 kb total) were excluded from the local window set to avoid confounding signals from TE-altered spatial interaction. This normalization successfully corrected the systemic bias between samples (Figure S8).

To compare the normalized IF value between homologous alleles, we estimated log2 fold change as the log2 ratio of the mean IF values of the TE-containing allele relative to that of the homologous TE-free alleles. We also estimated z-score, which is the difference in the mean IF values between TE-containing and TE-free alleles, normalized by the standard deviation across replicates and samples.

### CUT&Tag data analysis

The A4 and A7 CUT&Tag data targeting H3K9me3 were obtained from ((Huang et al. 2025); BioProject PRJNA1159377). We processed data following the CUT&Tag Data Processing and Analysis Tutorial (Zheng et al. 2020; Henikoff et al. 2020), estimating the histone-modification magnitude (HM) at each position based on sequencing coverage. To assess H3K9me3 enrichment around TEs, we calculated the mean HM for 5 kb flanking regions (upstream and downstream) of each TE and normalized this against the mean HM of local background windows (20–40 kb upstream and downstream), following (Lee and Karpen 2017). Normalized HM values >1 were considered H3K9me3-enriched.

### Identification of EU/PCH boundaries

CUT&Tag data targeting H3K9me3 were aligned to the genome assemblies of the corresponding genome assembly using bwa mem (v0.7.17-r1188), filtered using samtools (v 1.16) (Danecek et al. 2021) to retain reads of MAPQ > 30. Genome-wide H3K9me3 coverage tracks were generated in bedgraph format using bamCoverage (deepTools v3.5.1) (Ramírez et al. 2016) and visualized in the Integrative Genomics Viewer (IGV) (Thorvaldsdóttir et al. 2013) to assess H3K9me3 distribution along chromosome arms. EU/PCH boundaries were empirically defined based on H3K9me3 enrichment patterns (Figure S2). To ensure consistent boundary definitions across strains for downstream analyses, CUT&Tag reads were also aligned to the *D. melanogaster* iso-1 Release 6 reference genome. Empirical boundaries defined in this common coordinate system were lifted over to A4 and A7 genome coordinates and visually verified against strain-specific H3K9me3 enrichment profiles.

### RNA-sequencing experiment and analysis

Total RNA was extracted using the Zymo Direct-zol RNA Miniprep Kits. Library preparation and sequencing with Illumina NovaSeq X Plus were performed by NovoGene (Sacramento, CA). Each genotype has three biological replicates with an average of 30 million reads per sample. The *D. melanogaster* genome annotation file (FlyBase r6.59, dmel-all-r6.59.gtf) was obtained from FlyBase and lifted over from the iso-1 reference genome to the A4 and A7 assemblies using Liftoff (v1.6.3) (Shumate and Salzberg 2021). Raw RNA-seq reads were adapter-trimmed and quality-filtered using fastp (v0.23.4) (Chen et al. 2018) and then aligned to the genome assemblies of the corresponding genotype using STAR (v2.7.11b) (Dobin et al. 2013). Gene expression level was quantified using RSEM (v1.3.3) (Li and Dewey 2011), and the resulting FPKM values served as input for rank-based differential expression analyses.

### Conversion of TE coordinates between genomes

Some genomic attributes (e.g., chromatin state) analyzed were previously reported in coordinates based on *D. melanogaster* iso-1 reference genome. We converted the location of the A4 and A7 TEs to Release 6 coordinates by aligning the A4 or A7 genome to the Release 6 assembly using minimap2 (v2.24) (Li 2018), followed by the *call* command within *paftools.js* (part of the minimap2 package).

### Estimation of TE population frequencies

Population frequencies were estimated using Zambian population samples (BioProject SRP006733) from the Drosophila Genome Nexus resource (Lack et al. 2015), which represent the ancestral range of *Drosophila melanogaster*. We applied the computational pipeline previously developed in (Lee 2021; Huang et al. 2022) to estimate the frequencies of TEs identified in the A4 and A7 assemblies.

### Estimation of TE age

Multiple genome alignment was generated in reference-free mode using genome assemblies from this study (A4 and A7) and 12 DSPR genomes (Chakraborty et al. 2019; BioProject PRJNA418342). Structural variants were called, and TEs were identified using the Drosophila transposon canonical sequence library from the Bergman lab using Pannagram (Igolkina et al. 2025b, 2025c). We computed pairwise sequence similarity among TEs, defining two TEs as “similar” if they shared at least 85% alignment coverage with at least 85% sequence identity. In the resulting connection graph, “hubs” were defined as groups of at least 3 mutually similar TEs. Hubs were assigned to a TE family or superfamily if greater than 95% of constituent TEs belonged to that classification. Directed edges represented unidirectional similarities indicative of nested TEs, where greater than 85% of a query TE (arrow origin) aligned to a reference TE (arrow target), but not vice versa.

## Acknowledgement

We would like to thank Tony Long for providing the studied strains, J.J. Emerson for extensive discussion of the project, and Min-Chi Yang for technical assistance. We appreciate the University of California High-Throughput Genomics Facility and High-Performance Cluster at the University of California, Irvine for sequencing and computational resources. We also thank Aniek Janssen, Evgeny Kvon, and Brandon Gaut for critically reading the manuscript. This study was supported by NIH R35GM142494 to YCGL.

## References

Ahmed I, Sarazin A, Bowler C, Colot V, Quesneville H. 2011. Genome-wide evidence for local DNA methylation spreading from small RNA-targeted sequences in Arabidopsis. Nucleic Acids Res 39: 6919–6931.

B Charlesworth, Langley CH. 1989. The Population Genetics of Drosophila Transposable Elements. Annual Review of Genetics 23: 251–287.

Ballouz S, Dobin A, Gillis JA. 2019. Is it time to change the reference genome? Genome Biol 20: 159.

Barrón MG, Fiston-Lavier A-S, Petrov DA, González J. 2014. Population Genomics of Transposable Elements in Drosophila. Annual Review of Genetics 48: 561–581.

Becker JS, McCarthy RL, Sidoli S, Donahue G, Kaeding KE, He Z, Lin S, Garcia BA, Zaret KS. 2017. Genomic and Proteomic Resolution of Heterochromatin and Its Restriction of Alternate Fate Genes. Molecular Cell 68: 1023–1037.e15.

Bedford T, Hartl DL. 2009. Optimization of gene expression by natural selection. Proceedings of the National Academy of Sciences 106: 1133–1138.

Bellen HJ, Levis RW, Liao G, He Y, Carlson JW, Tsang G, Evans-Holm M, Hiesinger PR, Schulze KL, Rubin GM, et al. 2004. The BDGP Gene Disruption Project: Single Transposon Insertions Associated With 40% of Drosophila Genes. Genetics 167: 761– 781.

Bergman CM, Bensasson D. 2007. Recent LTR Retrotransposon Insertion Contrasts with Waves of Non-LTR Insertion Since Speciation in Drosophila Melanogaster. PNAS 104: 11340–11345.

Blumenstiel JP, Chen X, He M, Bergman CM. 2014. An Age-of-Allele Test of Neutrality for Transposable Element Insertions. Genetics 196: 523–538.

Bourque G, Burns KH, Gehring M, Gorbunova V, Seluanov A, Hammell M, Imbeault M, Izsvák Z, Levin HL, Macfarlan TS, et al. 2018. Ten things you should know about transposable elements. Genome Biology 19: 199.

Bystricky K, Merkenschlager M. 2020. Editorial overview: Genome architecture and expression. Current Opinion in Genetics & Development 61: iii–vi.

Chakraborty M, Emerson JJ, Macdonald SJ, Long AD. 2019. Structural variants exhibit widespread allelic heterogeneity and shape variation in complex traits. Nature Communications 10: 1–11.

Chandradoss KR, Guthikonda PK, Kethavath S, Dass M, Singh H, Nayak R, Kurukuti S, Sandhu KS. 2020. Biased visibility in Hi-C datasets marks dynamically regulated condensed and decondensed chromatin states genome-wide. BMC Genomics 21: 175.

Chen N-C, Solomon B, Mun T, Iyer S, Langmead B. 2021. Reference flow: reducing reference bias using multiple population genomes. Genome Biol 22: 8.

Chen S, Zhou Y, Chen Y, Gu J. 2018. fastp: an ultra-fast all-in-one FASTQ preprocessor. Bioinformatics 34: i884–i890.

Chen Z, Snetkova V, Bower G, Jacinto S, Clock B, Dizehchi A, Barozzi I, Mannion BJ, Alcaina-Caro A, Lopez-Rios J, et al. 2024. Increased enhancer–promoter interactions during developmental enhancer activation in mammals. Nat Genet 56: 675–685.

Cheng H, Concepcion GT, Feng X, Zhang H, Li H. 2021. Haplotype-resolved de novo assembly using phased assembly graphs with hifiasm. Nat Methods 18: 170–175.

Chiolo I, Minoda A, Colmenares SU, Polyzos A, Costes SV, Karpen GH. 2011. Double-Strand Breaks in Heterochromatin Move Outside of a Dynamic HP1a Domain to Complete Recombinational Repair. Cell 144: 732–744.

Choi JY, Lee YCG. 2020. Double-edged sword: The evolutionary consequences of the epigenetic silencing of transposable elements. PLOS Genetics 16: e1008872.

Choudhary MN, Friedman RZ, Wang JT, Jang HS, Zhuo X, Wang T. 2020. Co-opted transposons help perpetuate conserved higher-order chromosomal structures. Genome Biology 21: 16.

Choudhary MNK, Quaid K, Xing X, Schmidt H, Wang T. 2023. Widespread contribution of transposable elements to the rewiring of mammalian 3D genomes. Nature Communications 14: 634.

Chuong EB, Elde NC, Feschotte C. 2017. Regulatory activities of transposable elements: from conflicts to benefits. Nature Reviews Genetics 18: 71–86.

Clowney EJ, LeGros MA, Mosley CP, Clowney FG, Markenskoff-Papadimitriou EC, Myllys M, Barnea G, Larabell CA, Lomvardas S. 2012. Nuclear Aggregation of Olfactory Receptor Genes Governs Their Monogenic Expression. Cell 151: 724–737.

Coronado-Zamora M, González J. 2025. The epigenetics effects of transposable elements are genomic context dependent and not restricted to gene silencing in Drosophila. Genome Biol 26: 251.

Cosby RL, Judd J, Zhang R, Zhong A, Garry N, Pritham EJ, Feschotte C. 2021. Recurrent evolution of vertebrate transcription factors by transposase capture. Science 371: eabc6405.

Cridland JM, Macdonald SJ, Long AD, Thornton KR. 2013. Abundance and Distribution of Transposable Elements in Two Drosophila QTL Mapping Resources. Mol Biol Evol 30: 2311–2327.

Csink AK, Henikoff S. 1996. Genetic modification of heterochromatic association and nuclear organization in Drosophila. Nature 381: 529–531.

Czech B, Munafò M, Ciabrelli F, Eastwood EL, Fabry MH, Kneuss E, Hannon GJ. 2018. piRNA-Guided Genome Defense: From Biogenesis to Silencing. Annual Review of Genetics 52: 131–157.

Daborn PJ, Yen JL, Bogwitz MR, Le Goff G, Feil E, Jeffers S, Tijet N, Perry T, Heckel D, Batterham P, et al. 2002. A Single P450 Allele Associated with Insecticide Resistance in Drosophila. Science 297: 2253–2256.

Danecek P, Bonfield JK, Liddle J, Marshall J, Ohan V, Pollard MO, Whitwham A, Keane T, McCarthy SA, Davies RM, et al. 2021. Twelve years of SAMtools and BCFtools. Gigascience 10: giab008.

Dekker J, Mirny LA. 2024. The chromosome folding problem and how cells solve it. Cell 187: 6424–6450.

Dekker J, Rippe K, Dekker M, Kleckner N. 2002. Capturing Chromosome Conformation. Science 295: 1306–1311.

Dernburg AF, Broman KW, Fung JC, Marshall WF, Philips J, Agard DA, Sedat JW. 1996. Perturbation of Nuclear Architecture by Long-Distance Chromosome Interactions. Cell 85: 745–759.

Diehl AG, Ouyang N, Boyle AP. 2020. Transposable elements contribute to cell and species-specific chromatin looping and gene regulation in mammalian genomes. Nature Communications 11: 1796.

Dobin A, Davis CA, Schlesinger F, Drenkow J, Zaleski C, Jha S, Batut P, Chaisson M, Gingeras TR. 2013. STAR: ultrafast universal RNA-seq aligner. Bioinformatics 29: 15–21.

Eichten SR, Ellis NA, Makarevitch I, Yeh C-T, Gent JI, Guo L, McGinnis KM, Zhang X, Schnable PS, Vaughn MW, et al. 2012. Spreading of Heterochromatin Is Limited to Specific Families of Maize Retrotransposons. PLOS Genet 8: e1003127.

Elliott TA, Gregory TR. 2015. Do larger genomes contain more diverse transposable elements? BMC Evolutionary Biology 15: 69.

Eres IE, Luo K, Hsiao CJ, Blake LE, Gilad Y. 2019. Reorganization of 3D genome structure may contribute to gene regulatory evolution in primates. PLOS Genetics 15: e1008278.

Falk M, Feodorova Y, Naumova N, Imakaev M, Lajoie BR, Leonhardt H, Joffe B, Dekker J, Fudenberg G, Solovei I, et al. 2019. Heterochromatin drives compartmentalization of inverted and conventional nuclei. Nature 570: 395–399.

Feschotte C. 2008. Transposable elements and the evolution of regulatory networks. Nat Rev Genet 9: 397–405.

Filion GJ, van Bemmel JG, Braunschweig U, Talhout W, Kind J, Ward LD, Brugman W, de Castro Genebra de Jesus I, Kerkhoven RM, Bussemaker HJ, et al. 2010. Systematic protein location mapping reveals five principal chromatin types in Drosophila cells. Cell 143: 212–224.

Fueyo R, Judd J, Feschotte C, Wysocka J. 2022. Roles of transposable elements in the regulation of mammalian transcription. Nat Rev Mol Cell Biol 1–17.

Gilbertson EN, Brand CM, McArthur E, Rinker DC, Kuang S, Pollard KS, Capra JA. 2024. Machine Learning Reveals the Diversity of Human 3D Chromatin Contact Patterns. Mol Biol Evol 41: msae209.

Giles KA, Taberlay PC, Cesare AJ, Jones MJK. 2025. Roles for the 3D genome in the cell cycle, DNA replication, and double strand break repair. Front Cell Dev Biol 13: 1548946.

Goel M, Sun H, Jiao W-B, Schneeberger K. 2019. SyRI: finding genomic rearrangements and local sequence differences from whole-genome assemblies. Genome Biol 20: 277.

Golov AK, Gavrilov AA, Kaplan N, Razin SV. 2024. A genome-wide nucleosome-resolution map of promoter-centered interactions in human cells corroborates the enhancer-promoter looping model. eLife 12. https://elifesciences.org/reviewed-preprints/91596 (Accessed October 30, 2025).

González J, Macpherson JM, Petrov DA. 2009. A Recent Adaptive Transposable Element Insertion Near Highly Conserved Developmental Loci in Drosophila melanogaster. Mol Biol Evol 26: 1949–1961.

Groen SC, Ćalić I, Joly-Lopez Z, Platts AE, Choi JY, Natividad M, Dorph K, Mauck WM, Bracken B, Cabral CLU, et al. 2020. The strength and pattern of natural selection on gene expression in rice. Nature 578: 572–576.

Guio L, Barrón MG, González J. 2014. The transposable element Bari-Jheh mediates oxidative stress response in Drosophila. Molecular Ecology 23: 2020–2030.

Harmon B, Sedat J. 2005. Cell-by-Cell Dissection of Gene Expression and Chromosomal Interactions Reveals Consequences of Nuclear Reorganization. PLOS Biology 3: e67.

Haynes KA, Gracheva E, Elgin SCR. 2007. A Distinct Type of Heterochromatin Within Drosophila melanogaster Chromosome 4. Genetics 175: 1539–1542.

Henikoff S, Henikoff JG, Kaya-Okur HS, Ahmad K. 2020. Efficient chromatin accessibility mapping in situ by nucleosome-tethered tagmentation eds. R. Bonasio, J.K. Tyler, C.G. Danko, and J. Cao. eLife 9: e63274.

Hoencamp C, Dudchenko O, Elbatsh AMO, Brahmachari S, Raaijmakers JA, van Schaik T, Sedeño Cacciatore Á, Contessoto VG, van Heesbeen RGHP, van den Broek B, et al. 2021. 3D genomics across the tree of life reveals condensin II as a determinant of architecture type. Science 372: 984–989.

Hof AE van’t, Campagne P, Rigden DJ, Yung CJ, Lingley J, Quail MA, Hall N, Darby AC, Saccheri IJ. 2016. The industrial melanism mutation in British peppered moths is a transposable element. Nature 534: 102–105.

Hoskins RA, Carlson JW, Wan KH, Park S, Mendez I, Galle SE, Booth BW, Pfeiffer BD, George RA, Svirskas R, et al. 2015. The Release 6 reference sequence of the Drosophila melanogaster genome. Genome Res gr.185579.114.

Huang Y, Gao ZY, Ly K, Lin L, Lambooij J-P, King EG, Janssen A, Wei KH-C, Lee YCG. 2025. Polymorphic transposable elements contribute to variation in recombination landscapes. Proceedings of the National Academy of Sciences 122: e2427312122.

Huang Y, Shukla H, Lee YCG. 2022. Species-specific chromatin landscape determines how transposable elements shape genome evolution eds. M. Nordborg and M. Przeworski. eLife 11: e81567.

Ibrahim DM, Mundlos S. 2020. The role of 3D chromatin domains in gene regulation: a multi-facetted view on genome organization. Current Opinion in Genetics & Development 61: 1–8.

Igolkina AA, Bezlepsky AD, Nordborg M. 2025a. Pannagram: unbiased pangenome alignment and the Mobilome calling. 2025.02.07.637071. https://www.biorxiv.org/content/10.1101/2025.02.07.637071v1 (Accessed May 5, 2025).

Igolkina AA, Bezlepsky AD, Nordborg M. 2025b. Pannagram: unbiased pangenome alignment and the Mobilome calling. 2025.02.07.637071. https://www.biorxiv.org/content/10.1101/2025.02.07.637071v1 (Accessed December 24, 2025).

Igolkina AA, Vorbrugg S, Rabanal FA, Liu H-J, Ashkenazy H, Kornienko AE, Fitz J, Collenberg M, Kubica C, Mollá Morales A, et al. 2025c. A comparison of 27 Arabidopsis thaliana genomes and the path toward an unbiased characterization of genetic polymorphism. Nat Genet 57: 2289–2301.

James TC, Elgin SC. 1986. Identification of a nonhistone chromosomal protein associated with heterochromatin in Drosophila melanogaster and its gene. Molecular and Cellular Biology 6: 3862–3872.

Janssen A, Breuer GA, Brinkman EK, Meulen AI van der, Borden SV, Steensel B van, Bindra RS, LaRocque JR, Karpen GH. 2016. A single double-strand break system reveals repair dynamics and mechanisms in heterochromatin and euchromatin. Genes Dev 30: 1645–1657.

Janssen A, Colmenares SU, Karpen GH. 2018. Heterochromatin: Guardian of the Genome. Annual Review of Cell and Developmental Biology 34: 265–288.

Kaya-Okur HS, Wu SJ, Codomo CA, Pledger ES, Bryson TD, Henikoff JG, Ahmad K, Henikoff S. 2019. CUT&Tag for efficient epigenomic profiling of small samples and single cells. Nature Communications 10: 1930.

Keenen MM, Brown D, Brennan LD, Renger R, Khoo H, Carlson CR, Huang B, Grill SW, Narlikar GJ, Redding S. 2021. HP1 proteins compact DNA into mechanically and positionally stable phase separated domains eds. S. Deindl and K. Struhl. eLife 10: e64563.

Kelleher ES, Barbash DA, Blumenstiel JP. 2020. Taming the Turmoil Within: New Insights on the Containment of Transposable Elements. Trends in Genetics 36: 474–489.

King EG, Macdonald SJ, Long AD. 2012. Properties and Power of the Drosophila Synthetic Population Resource for the Routine Dissection of Complex Traits. Genetics 191: 935– 949.

Klein KN, Zhao PA, Lyu X, Sasaki T, Bartlett DA, Singh AM, Tasan I, Zhang M, Watts LP, Hiraga S, et al. 2021. Replication timing maintains the global epigenetic state in human cells. Science 372: 371–378.

Kragesteen BK, Spielmann M, Paliou C, Heinrich V, Schöpflin R, Esposito A, Annunziatella C, Bianco S, Chiariello AM, Jerković I, et al. 2018. Dynamic 3D chromatin architecture contributes to enhancer specificity and limb morphogenesis. Nat Genet 50: 1463–1473.

Lack JB, Cardeno CM, Crepeau MW, Taylor W, Corbett-Detig RB, Stevens KA, Langley CH, Pool JE. 2015. The Drosophila Genome Nexus: A Population Genomic Resource of 623 Drosophila melanogaster Genomes, Including 197 from a Single Ancestral Range Population. Genetics genetics.115.174664.

Langley CH, Montgomery E, Hudson R, Kaplan N, Charlesworth B. 1988. On the role of unequal exchange in the containment of transposable element copy number. Genet Res 52: 223–235.

Langley SA, Miga KH, Karpen GH, Langley CH. 2019. Haplotypes spanning centromeric regions reveal persistence of large blocks of archaic DNA eds. M. Nordborg, D. Tautz, M. Nordborg, and A.G. Clark. eLife 8: e42989.

Larson AG, Elnatan D, Keenen MM, Trnka MJ, Johnston JB, Burlingame AL, Agard DA, Redding S, Narlikar GJ. 2017. Liquid droplet formation by HP1α suggests a role for phase separation in heterochromatin. Nature 547: 236–240.

Lawson HA, Liang Y, Wang T. 2023. Transposable elements in mammalian chromatin organization. Nat Rev Genet 1–12.

Lee YCG. 2021. Synergistic epistasis of the deleterious effects of transposable elements ed. T. Slotte. Genetics iyab211.

Lee YCG. 2015. The Role of piRNA-Mediated Epigenetic Silencing in the Population Dynamics of Transposable Elements in Drosophila melanogaster. PLoS Genet 11: e1005269.

Lee YCG, Karpen GH. 2017. Pervasive epigenetic effects of Drosophila euchromatic transposable elements impact their evolution. eLife 6. https://www.ncbi.nlm.nih.gov/pmc/articles/PMC5505702/.

Lee YCG, Langley CH. 2010. Transposable elements in natural populations of Drosophila melanogaster. Phil Trans R Soc B 365: 1219–1228.

Lee YCG, Ogiyama Y, Martins NMC, Beliveau BJ, Acevedo D, Wu C-ting, Cavalli G, Karpen GH. 2020. Pericentromeric heterochromatin is hierarchically organized and spatially contacts H3K9me2 islands in euchromatin. PLOS Genetics 16: e1008673.

Li B, Dewey CN. 2011. RSEM: accurate transcript quantification from RNA-Seq data with or without a reference genome. BMC Bioinformatics 12: 323.

Li C, Bonder MJ, Syed S, Jensen M, Consortium (HGSVC) HGSV, Group HFAW, Gerstein MB, Zody MC, Chaisson MJP, Talkowski ME, et al. 2024. An integrative TAD catalog in lymphoblastoid cell lines discloses the functional impact of deletions and insertions in human genomes. Genome Res 34: 2304–2318.

Li H. 2013. Aligning sequence reads, clone sequences and assembly contigs with BWA-MEM. http://arxiv.org/abs/1303.3997 (Accessed December 31, 2025).

Li H. 2018. Minimap2: pairwise alignment for nucleotide sequences. Bioinformatics 34: 3094– 3100.

Li H. 2021. New strategies to improve minimap2 alignment accuracy. Bioinformatics 37: 4572– 4574.

Li Y, Hu M, Shen Y. 2018. Gene regulation in the 3D genome. Human Molecular Genetics 27: R228–R233.

Lieberman-Aiden E, Berkum NL van, Williams L, Imakaev M, Ragoczy T, Telling A, Amit I, Lajoie BR, Sabo PJ, Dorschner MO, et al. 2009. Comprehensive Mapping of Long-Range Interactions Reveals Folding Principles of the Human Genome. Science 326: 289–293.

Lin L, Huang Y, McIntyre J, Chang C-H, Colmenares S, Lee YCG. 2024a. Prevalent Fast Evolution of Genes Involved in Heterochromatin Functions. Molecular Biology and Evolution 41: msae181.

Lin M-J, Iyer S, Chen N-C, Langmead B. 2024b. Measuring, visualizing, and diagnosing reference bias with biastools. Genome Biol 25: 101.

Magklara A, Yen A, Colquitt BM, Clowney EJ, Allen W, Markenscoff-Papadimitriou E, Evans ZA, Kheradpour P, Mountoufaris G, Carey C, et al. 2011. An epigenetic signature for monoallelic olfactory receptor expression. Cell 145: 555–570.

Maksakova IA, Romanish MT, Gagnier L, Dunn CA, van de Lagemaat LN, Mager DL. 2006. Retroviral Elements and Their Hosts: Insertional Mutagenesis in the Mouse Germ Line. PLoS Genet 2: e2.

Marco-Sola S, Moure JC, Moreto M, Espinosa A. 2021. Fast gap-affine pairwise alignment using the wavefront algorithm. Bioinformatics 37: 456–463.

McArthur E, Rinker DC, Cheng Y, Wang Q, Wang J, Gilbertson EN, Fudenberg G, Pittman M, Keough K, Yue F, et al. 2025. Reconstructing the 3D genome organization of Neanderthals reveals that chromatin folding shaped phenotypic and sequence divergence. bioRxiv 2022.02.07.479462.

Montgomery E, Charlesworth B, Langley CH. 1987. A test for the role of natural selection in the stabilization of transposable element copy number in a population of Drosophila melanogaster. Genet Res 49: 31–41.

Nagano T, Lubling Y, Várnai C, Dudley C, Leung W, Baran Y, Mendelson Cohen N, Wingett S, Fraser P, Tanay A. 2017. Cell-cycle dynamics of chromosomal organization at single-cell resolution. Nature 547: 61–67.

Naumova N, Imakaev M, Fudenberg G, Zhan Y, Lajoie BR, Mirny LA, Dekker J. 2013. Organization of the Mitotic Chromosome. Science 342: 948–953.

Norton HK, Phillips-Cremins JE. 2017. Crossed wires: 3D genome misfolding in human disease. J Cell Biol 216: 3441–3452.

Open2C, Abdennur N, Fudenberg G, Flyamer IM, Galitsyna AA, Goloborodko A, Imakaev M, Venev SV. 2024. Pairtools: From sequencing data to chromosome contacts. PLOS Computational Biology 20: e1012164.

Osmanski AB, Paulat NS, Korstian J, Grimshaw JR, Halsey M, Sullivan KAM, Moreno-Santillán DD, Crookshanks C, Roberts J, Garcia C, et al. 2023. Insights into mammalian TE diversity through the curation of 248 genome assemblies. Science 380: eabn1430.

Pasquesi GIM, Allen H, Ivancevic A, Barbachano-Guerrero A, Joyner O, Guo K, Simpson DM, Gapin K, Horton I, Nguyen LL, et al. 2024. Regulation of human interferon signaling by transposon exonization. Cell 187: 7621–7636.e19.

Penagos-Puig A, Furlan-Magaril M. 2020. Heterochromatin as an Important Driver of Genome Organization. Front Cell Dev Biol 8: 579137.

Quadrana L, Silveira AB, Mayhew GF, LeBlanc C, Martienssen RA, Jeddeloh JA, Colot V. 2016. The Arabidopsis thaliana mobilome and its impact at the species level. eLife 5: e15716.

Ramírez F, Bhardwaj V, Arrigoni L, Lam KC, Grüning BA, Villaveces J, Habermann B, Akhtar A, Manke T. 2018. High-resolution TADs reveal DNA sequences underlying genome organization in flies. Nat Commun 9: 189.

Ramírez F, Ryan DP, Grüning B, Bhardwaj V, Kilpert F, Richter AS, Heyne S, Dündar F, Manke T. 2016. deepTools2: a next generation web server for deep-sequencing data analysis. Nucleic Acids Res 44: W160–W165.

Rebollo R, Karimi MM, Bilenky M, Gagnier L, Miceli-Royer K, Zhang Y, Goyal P, Keane TM, Jones S, Hirst M, et al. 2011. Retrotransposon-Induced Heterochromatin Spreading in the Mouse Revealed by Insertional Polymorphisms. PLoS Genet 7: e1002301.

Rech GE, Radío S, Guirao-Rico S, Aguilera L, Horvath V, Green L, Lindstadt H, Jamilloux V, Quesneville H, González J. 2022. Population-scale long-read sequencing uncovers transposable elements associated with gene expression variation and adaptive signatures in Drosophila. Nat Commun 13: 1948.

Riddle NC, Elgin SCR. 2018. The Drosophila Dot Chromosome: Where Genes Flourish Amidst Repeats. Genetics 210: 757–772.

Riddle NC, Jung YL, Gu T, Alekseyenko AA, Asker D, Gui H, Kharchenko PV, Minoda A, Plachetka A, Schwartz YB, et al. 2012. Enrichment of HP1a on Drosophila Chromosome 4 Genes Creates an Alternate Chromatin Structure Critical for Regulation in this Heterochromatic Domain. PLOS Genetics 8: e1002954.

Riddle NC, Minoda A, Kharchenko PV, Alekseyenko AA, Schwartz YB, Tolstorukov MY, Gorchakov AA, Jaffe JD, Kennedy C, Linder-Basso D, et al. 2011. Plasticity in patterns of histone modifications and chromosomal proteins in Drosophila heterochromatin. Genome Res 21: 147–163.

Ryba T, Hiratani I, Lu J, Itoh M, Kulik M, Zhang J, Schulz TC, Robins AJ, Dalton S, Gilbert DM. 2010. Evolutionarily conserved replication timing profiles predict long-range chromatin interactions and distinguish closely related cell types. Genome Res 20: 761–770.

Schmidt D, Schwalie PC, Wilson MD, Ballester B, Gonçalves Â, Kutter C, Brown GD, Marshall A, Flicek P, Odom DT. 2012. Waves of Retrotransposon Expansion Remodel Genome Organization and CTCF Binding in Multiple Mammalian Lineages. Cell 148: 335–348.

Sexton T, Yaffe E, Kenigsberg E, Bantignies F, Leblanc B, Hoichman M, Parrinello H, Tanay A, Cavalli G. 2012. Three-Dimensional Folding and Functional Organization Principles of the Drosophila Genome. Cell 148: 458–472.

Shanta O, Noor A, Chaisson MJP, Sanders AD, Zhao X, Malhotra A, Porubsky D, Rausch T, Gardner EJ, Rodriguez OL, et al. 2020. The effects of common structural variants on 3D chromatin structure. BMC Genomics 21: 95.

Shukla HG, Bawa PS, Srinivasan S. 2019. hg19KIndel: ethnicity normalized human reference genome. BMC Genomics 20: 459.

Shukla HG, Chakraborty M, Emerson JJ. 2025. Genetic variation in recalcitrant repetitive regions of the Drosophila melanogaster genome. Genome Res 35: 2023–2040.

Shumate A, Salzberg SL. 2021. Liftoff: accurate mapping of gene annotations. Bioinformatics 37: 1639–1643.

Slotkin RK, Martienssen R. 2007. Transposable elements and the epigenetic regulation of the genome. Nat Rev Genet 8: 272–285.

Smit AFA, Hubley R, Green P. 2015. RepeatMasker Open-4.0. 2013–2015.

Spielmann M, Lupiáñez DG, Mundlos S. 2018. Structural variation in the 3D genome. Nature Reviews Genetics 19: 453–467.

Stewart C, Kural D, Strömberg MP, Walker JA, Konkel MK, Stütz AM, Urban AE, Grubert F, Lam HYK, Lee W-P, et al. 2011. A Comprehensive Map of Mobile Element Insertion Polymorphisms in Humans. PLoS Genet 7: e1002236.

Strom AR, Emelyanov AV, Mir M, Fyodorov DV, Darzacq X, Karpen GH. 2017. Phase separation drives heterochromatin domain formation. Nature 547: 241–245.

Stuart T, Eichten SR, Cahn J, Karpievitch YV, Borevitz JO, Lister R. 2016. Population scale mapping of transposable element diversity reveals links to gene regulation and epigenomic variation. eLife 5: e20777.

Stutzman AV, Hill CA, Armstrong RL, Gohil R, Duronio RJ, Dowen JM, McKay DJ. 2024. Heterochromatic 3D genome organization is directed by HP1a- and H3K9-dependent and independent mechanisms. Molecular Cell 84: 2017–2035.e6.

Sudmant PH, Rausch T, Gardner EJ, Handsaker RE, Abyzov A, Huddleston J, Zhang Y, Ye K, Jun G, Fritz MH-Y, et al. 2015. An integrated map of structural variation in 2,504 human genomes. Nature 526: 75–81.

Symer DE, Connelly C, Szak ST, Caputo EM, Cost GJ, Parmigiani G, Boeke JD. 2002. Human L1 Retrotransposition Is Associated with Genetic Instability In Vivo. Cell 110: 327–338.

Thorburn D-MJ, Sagonas K, Binzer-Panchal M, Chain FJJ, Feulner PGD, Bornberg-Bauer E, Reusch TBH, Samonte-Padilla IE, Milinski M, Lenz TL, et al. 2023. Origin matters: Using a local reference genome improves measures in population genomics. Mol Ecol Resour 23: 1706–1723.

Thorvaldsdóttir H, Robinson JT, Mesirov JP. 2013. Integrative Genomics Viewer (IGV): high-performance genomics data visualization and exploration. Brief Bioinform 14: 178–192.

Thybert D, Roller M, Navarro FCP, Fiddes I, Streeter I, Feig C, Martin-Galvez D, Kolmogorov M, Janoušek V, Akanni W, et al. 2018. Repeat associated mechanisms of genome evolution and function revealed by the Mus caroli and Mus pahari genomes. Genome Res 28: 448–459.

Torosin NS, Anand A, Golla TR, Cao W, Ellison CE. 2020. 3D genome evolution and reorganization in the Drosophila melanogaster species group. PLOS Genetics 16: e1009229.

Torosin NS, Golla TR, Lawlor MA, Cao W, Ellison CE. 2022. Mode and Tempo of 3D Genome Evolution in Drosophila. Mol Biol Evol 39: msac216.

Wells JN, Feschotte C. 2020. A Field Guide to Eukaryotic Transposable Elements. Annual Review of Genetics 54: 539–561.

Wenger AM, Peluso P, Rowell WJ, Chang P-C, Hall RJ, Concepcion GT, Ebler J, Fungtammasan A, Kolesnikov A, Olson ND, et al. 2019. Accurate circular consensus long-read sequencing improves variant detection and assembly of a human genome. Nat Biotechnol 37: 1155–1162.

Wu X, Xiong D, Liu R, Lai X, Tian Y, Xie Z, Chen L, Hu L, Duan J, Gao X, et al. 2025. Evolutionary divergence in CTCF-mediated chromatin topology drives transcriptional innovation in humans. Nat Commun 16: 2941.

Zenk F, Zhan Y, Kos P, Löser E, Atinbayeva N, Schächtle M, Tiana G, Giorgetti L, Iovino N. 2021. HP1 drives de novo 3D genome reorganization in early Drosophila embryos. Nature 593: 289–293.

Zhang H, Blobel GA. 2023. Genome folding dynamics during the M-to-G1-phase transition. Current Opinion in Genetics & Development 80: 102036.

Zheng Y, Ahmad K, Henikoff S. 2020. CUT&Tag Data Processing and Analysis Tutorial. https://www.protocols.io/view/cut-amp-tag-data-processing-and-analysis-tutorial-bjk2kkye (Accessed December 24, 2025).

