## Supplementary Figures for "Polymorphic 3D genome architecture mediated by transposable elements"

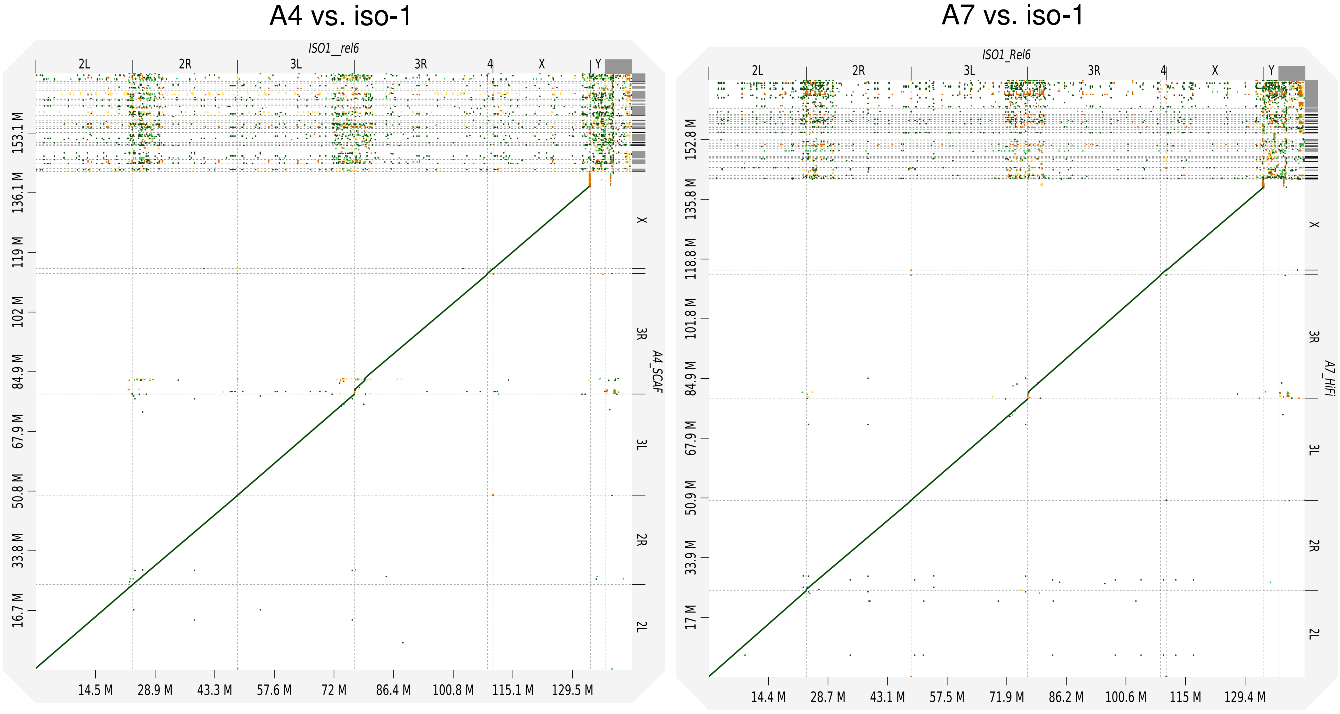


**Figure S1. Dot plots comparing PacBio HiFi *de novo* assemblies to the reference genome.** Dot plots illustrating synteny and alignment between the *D. melanogaster* iso-1 reference genome (x-axis) and the *de novo* PacBio HiFi assemblies (y-axis) for strain A4 (left) and strain A7 (right). The high linearity and lack of large-scale structural divergences confirm global structural integrity and high quality of the strain-specific PacBio assemblies.

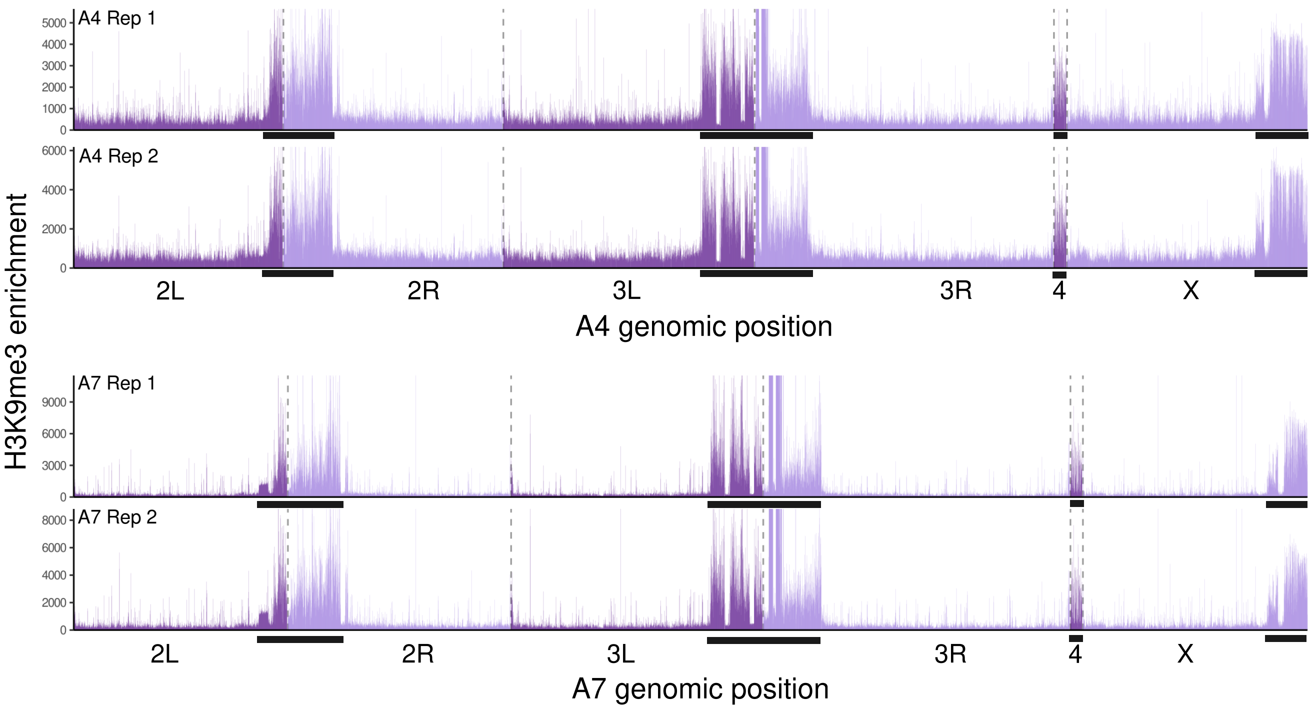


**Figure S2. Definition of pericentromeric heterochromatin (PCH) boundaries using genome-wide enrichment of H3K9me3.** H3K9me3 CUT&Tag coverage tracks for A4 (top panels; two replicates) and A7 (bottom panels; two replicates). Black bars below the tracks indicate the empirically defined PCH regions. Dashed vertical lines demarcate the arms/chromosomes. Note that Y-linked and repetitive unplaced contigs were also classified as PCH but are not displayed in this figure because they consist of multiple fragmented contigs.

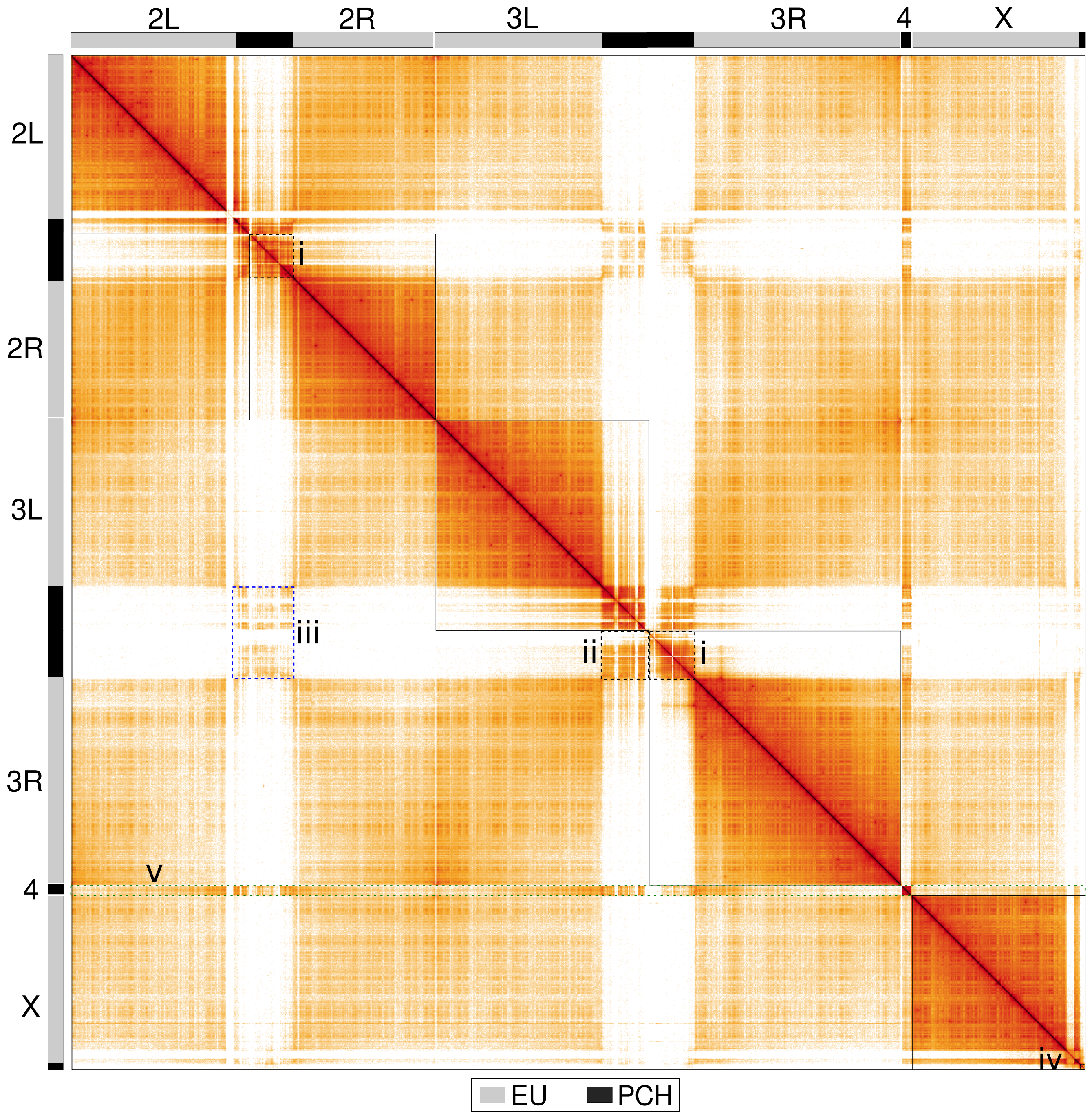


**Figure S3. Genome-wide Hi-C contact map for the A7 strain.** Solid boxes delineate chromosome territories. Highlighted key 3D structural features (in dashed line boxes): (i) PCH of each chromosomal arm forms distinct domains; (ii) more frequent interactions occur between PCH of arms from the same chromosome than between PCH of different chromosomes; (iii) spatial clustering of PCH across chromosomes; (iv) PCH of X chromosomes exhibits clear separation from its euchromatic arm; (v) the 4th chromosome interacts with both euchromatin and PCH of other chromosomes in similar frequencies

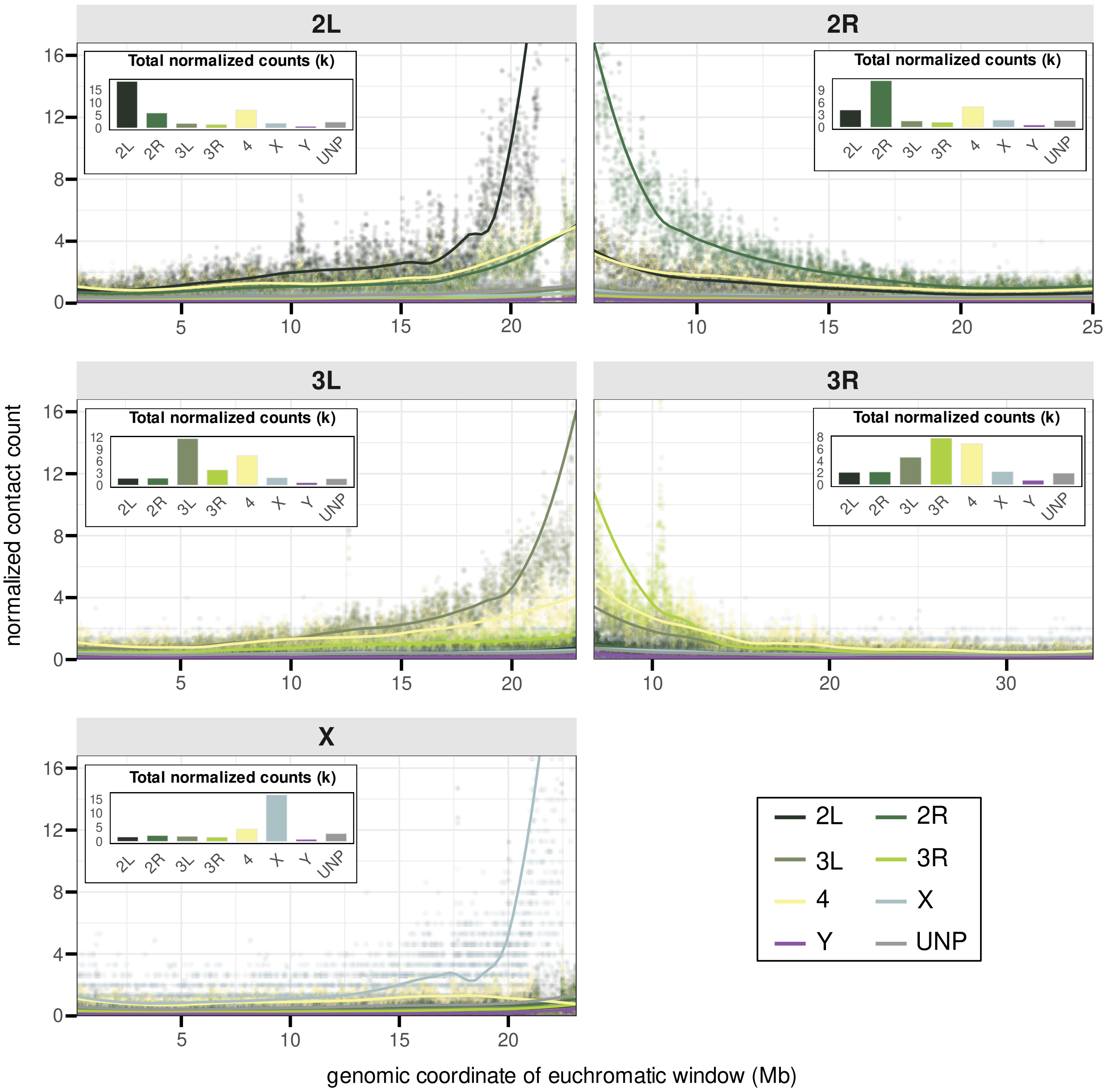


**Figure S4. Spatial interactions between euchromatin and PCH in A7 strain (Rep1).** Number of Hi-C reads supporting 3D interactions between 5kb euchromatic windows and PCH, normalized by the uniquely mappable size of PCH for 7. Lines/points of different colors represent interactions with PCH of a specific arm (2L, 2R, 3L, 3R, X, 4, Y) or unmapped (UNP). Insets show the total normalized contact counts for each arm. See Figure S5 and S6 for normalization using proportion of reads, which show consistent results.

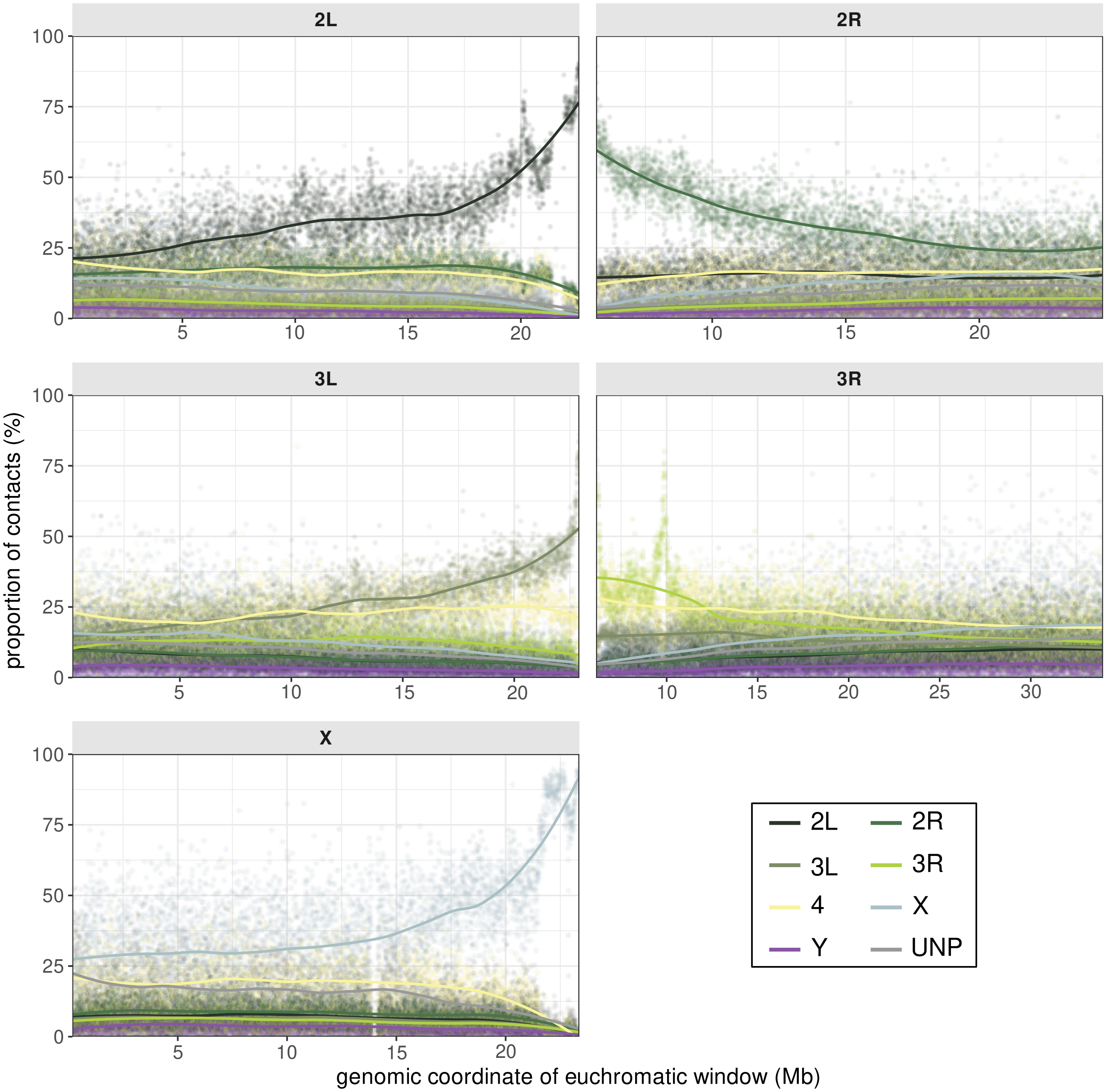


**Figure S5. Spatial interactions between euchromatin and PCH normalized as proportion in A4 strain.** The number of HiC-reads (from Fig 2B) supporting 3D interactions between 5kb euchromatic windows and PCH of a specific chromosome arm, out from all number of read pairs with one end mapped to the euchromatic window and other end mapped to the entirety of PCH.

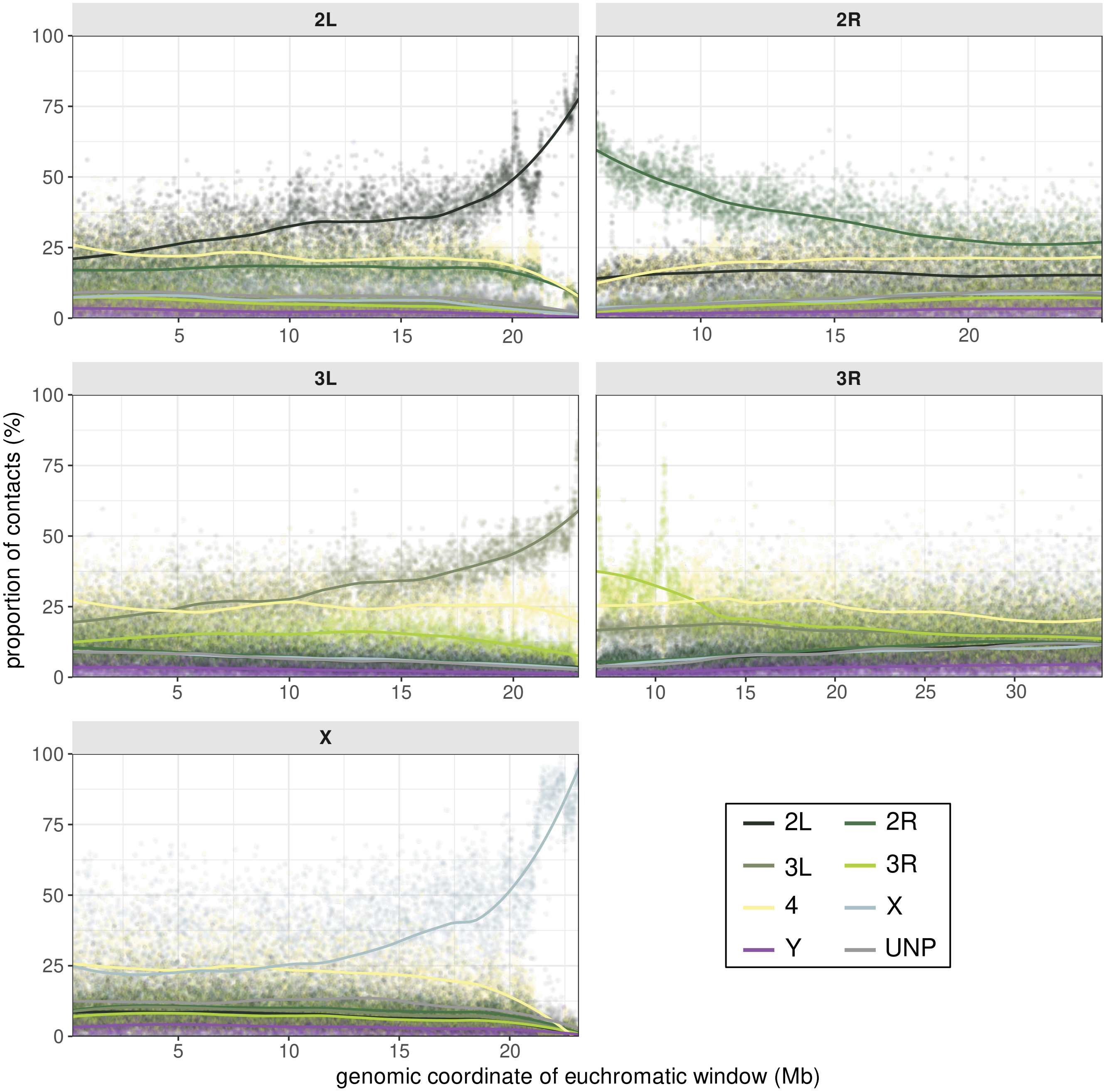


**Figure S6. Spatial interactions between euchromatin and PCH normalized as proportion in A7 strain.** The number of HiC-reads (from SFigure S4) supporting 3D interactions between 5kb euchromatic windows and PCH of a specific chromosome arm, out from all number of read pairs with one end mapped to the euchromatic window and other end mapped to the entirety of PCH

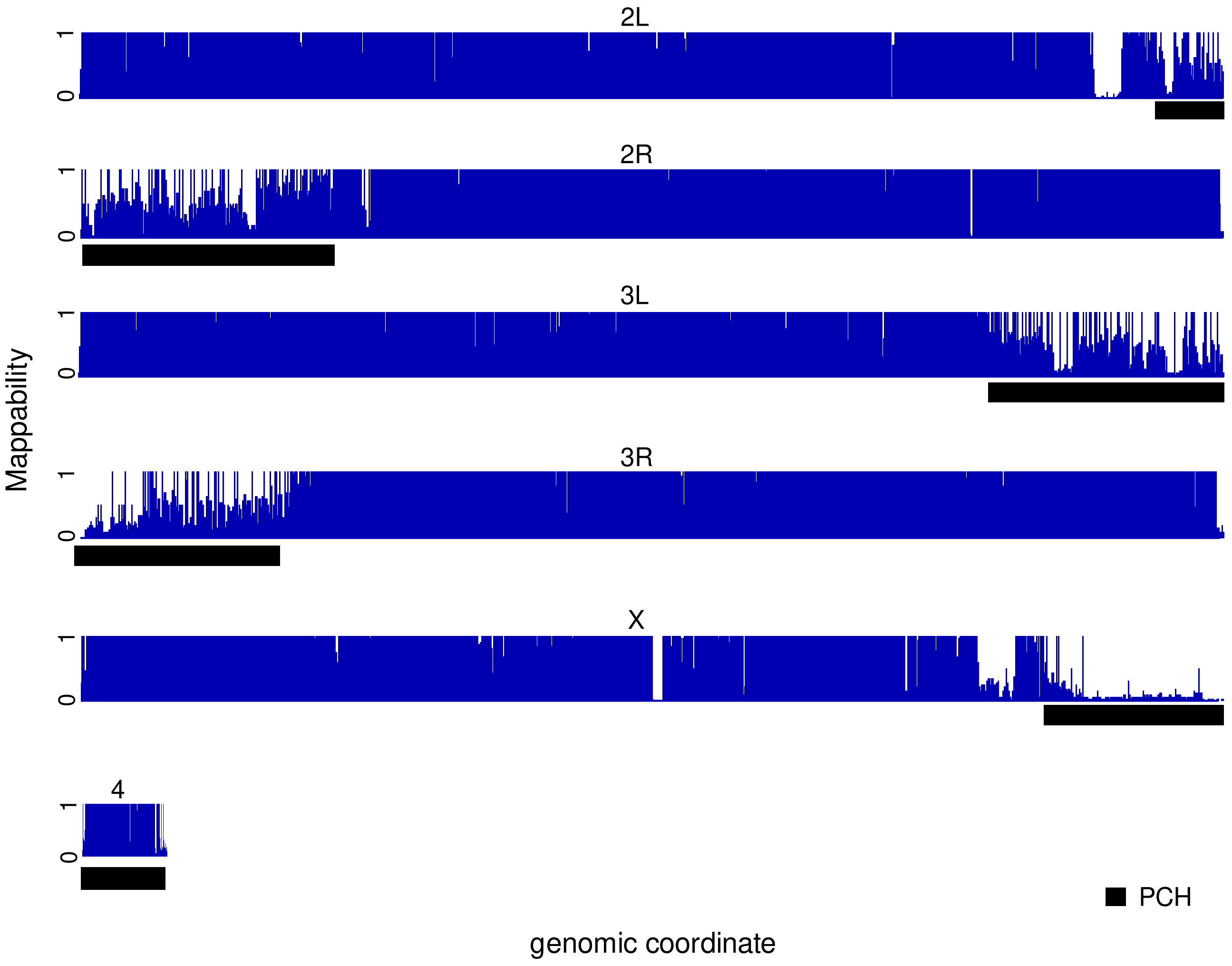


**Figure S7. Genome-wide mappability tracks from IGV.** Mappability scores (0–1) across the A4 genome calculated for k=150. PCH, as indicated by black bars, exhibit significantly reduced mappability due to high repetitive content, necessitating the "rescue" strategy for multi-mapping reads employed in this study.

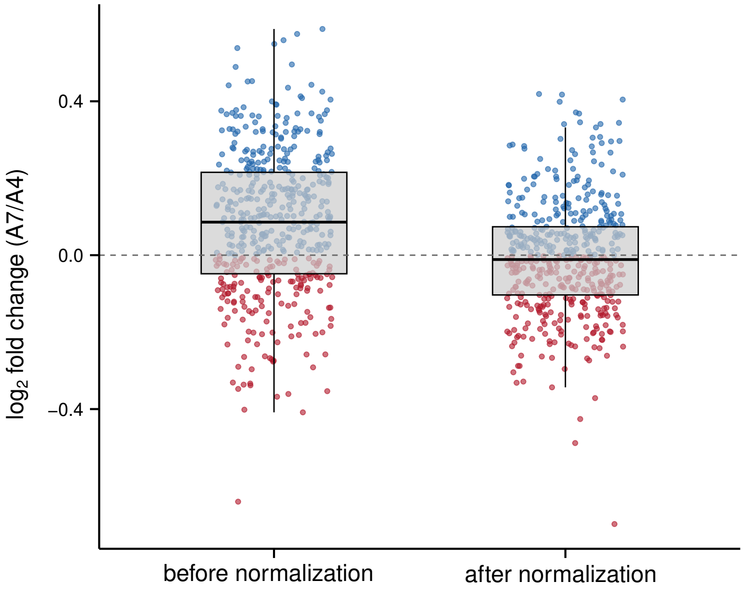


**Figure S8. Effectiveness of the normalization strategy.** Box plots showing the distribution of $\log_2$ fold changes ($\log_2$(A7/A4)) in interaction frequency for randomly selected TE-free windows before (left) and after (right) the proposed two-stage normalization. **Left:** Before normalization, log2 fold changes skew positively (median > 0), reflecting the higher library complexity (or more long-range interactions) of A7 (Table S1). **Right:** After applying the localized normalization correction, the distribution of $\log_2$ fold changes centers around zero, indicating the effective removal of technical bias

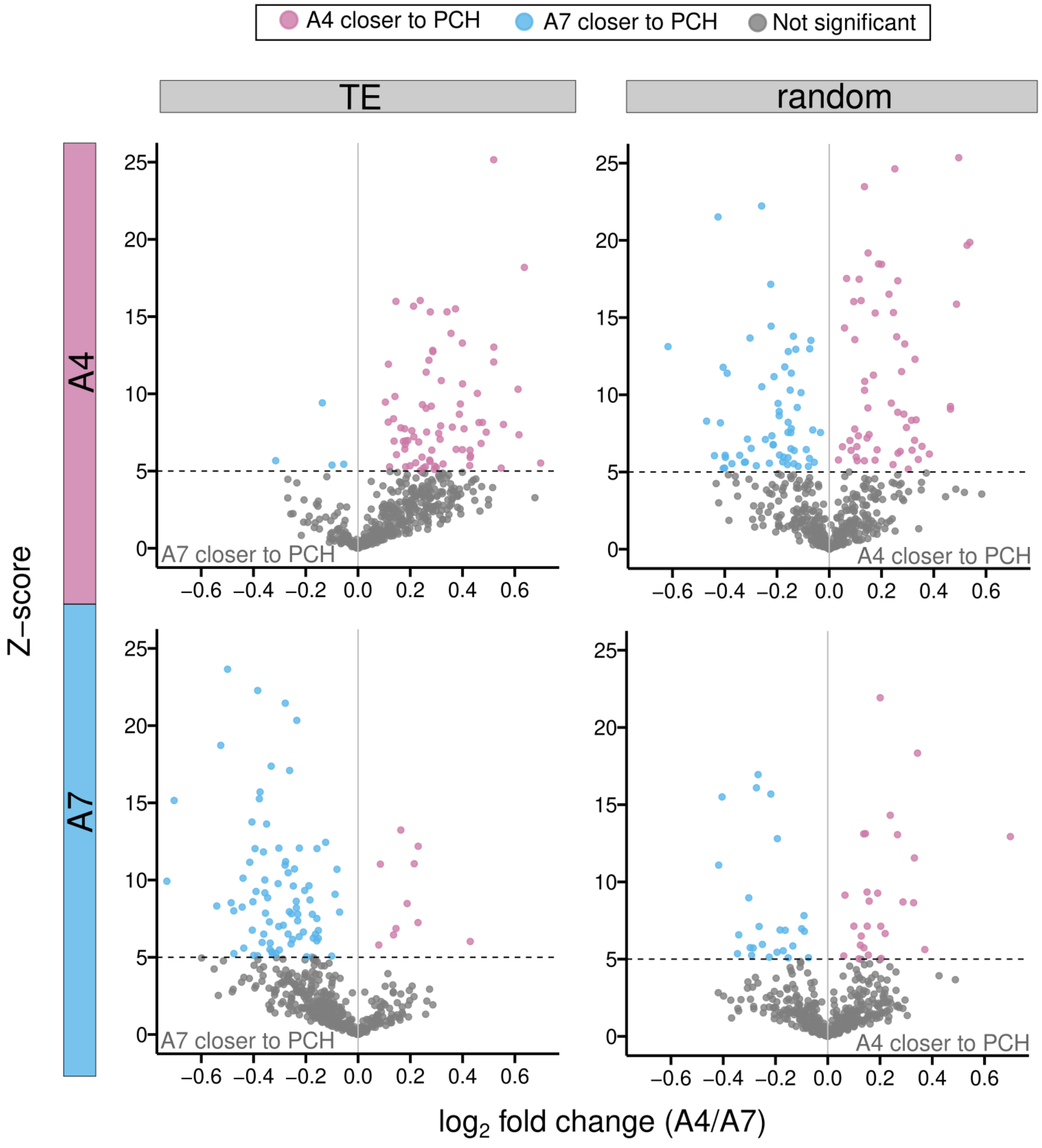


**Figure S9. Volcano plots comparing differential PCH interactions between homologous alleles using only unique-unique (UU) reads.** Volcano plots comparing differential PCH interaction between homologous alleles for TEs (left) and randomly selected TE-free windows (right) in A4 (top) and A7 (bottom), generated exclusively using UU Hi-C read pairs and without including rescued multi-mapping reads. Even with this reduced dataset, TE-containing alleles (left) exhibit a significant enrichment of PCH interactions compared to random controls (right), confirming that the observed TE-PCH 3D interaction is robust to the inclusion (Figure 2C) or exclusion of rescued multi-mapping reads.

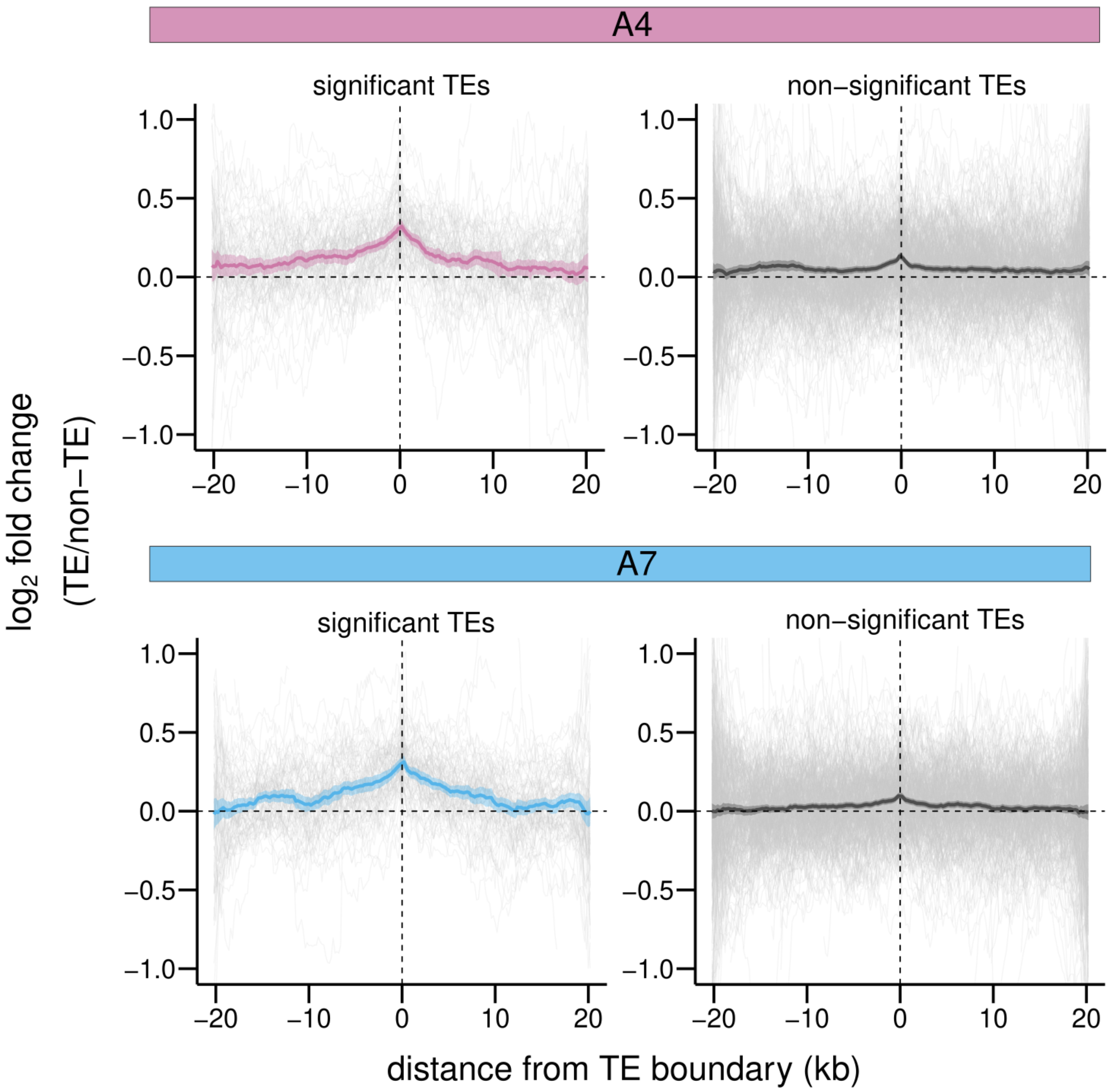


**Figure S10. Averaged $\log_2$ fold change of IF values between homologous with and without TE alleles using only unique-unique (UU) reads.** The log2 fold change of IF values between homologous with and without TE alleles averaged over TEs with and without significant interactions with PCH, generated exclusively using UU Hi-C read pairs and without including rescued multi-mapping reads. The observed spatial extent of TE-mediated PCH 3D proximity (~10 kb) is consistent with the results obtained including rescued multi-mapping reads (Figure 2E).

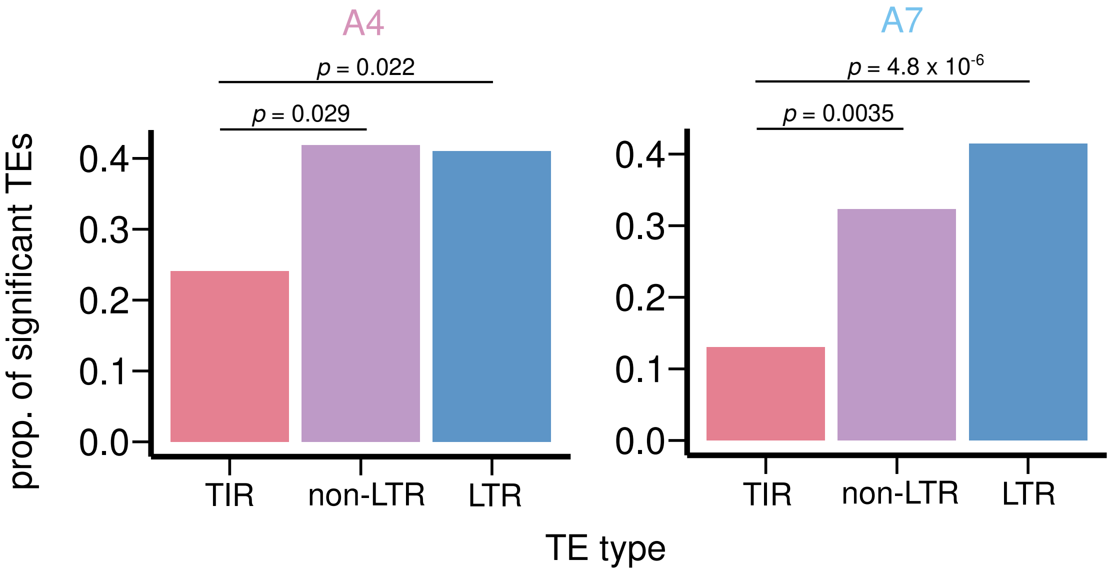


**Figure S11. Proportion of TEs with significant PCH 3D interactions for different TE type.** RNA-based TEs, LTR and non-LTR, show a higher proportion of significant PCH interactions compared to DNA-based (TIR) TEs in both strains (*Fisher's Exact test,* LTR *v.s.* TIR, odds ratio = 2.19, $p$ = 0.022; non-LTR *v.s.*. TIR, odds ratio = 2.27, $p$ = 0.029 (A4). LTR *v.s.* TIR, odds ratio = 4.71, $p = 4.8 \times 10^{-6}$; non-LTR *v.s.* TIR, odds ratio = 3.16, $p$ = 0.0035 (A7)).

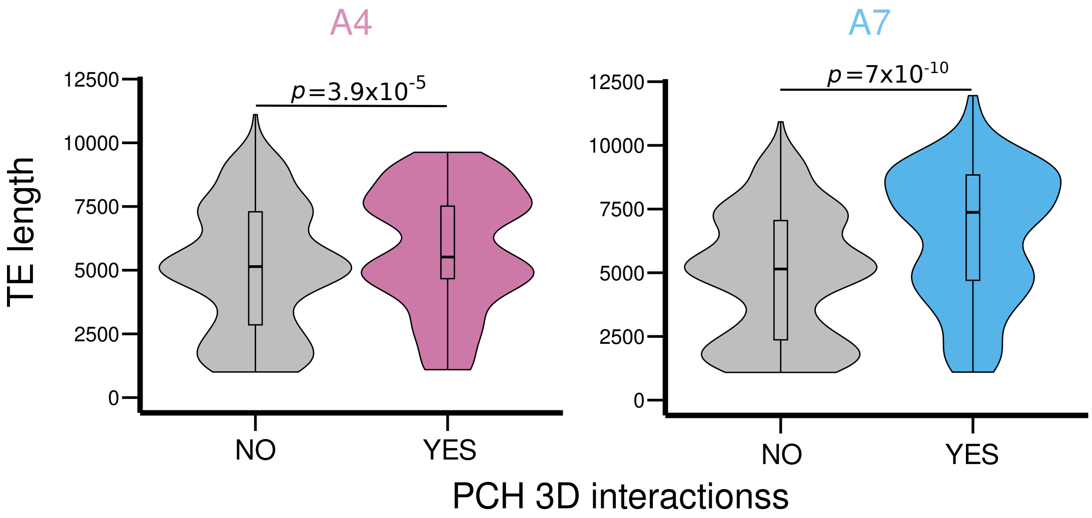


**Figure S12. Violin plots comparing length of TEs with and without significant PCH interactions.** In both strains, TEs that interact with PCH are significantly longer than those that do not (A4: median length, 6,996 bp (with) *v.s.* 5,144 bp (without), *Mann-Whitney U test, p* = $3.9 \times 10^{-5}$; A7: median length, 7,380 bp (with) *v.s.* 5,419 bp (without), *Mann-Whitney U test, p* = $7 \times 10^{-10}$).

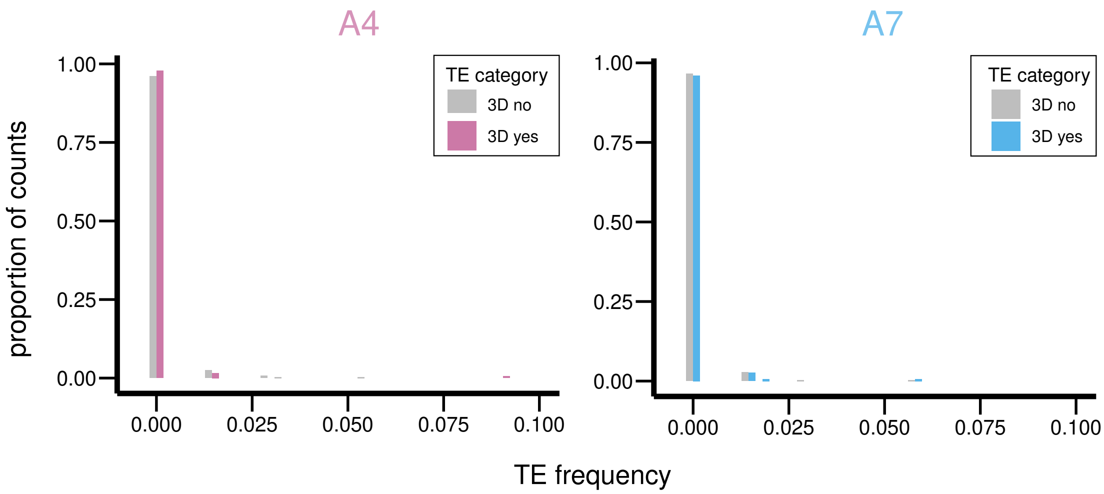


**Figure S13. Distribution of population frequencies of TEs with and without PCH 3D interactions.** Both categories of TEs are predominantly segregating at very low frequencies (< 0.05), limiting the statistical power to detect differences.

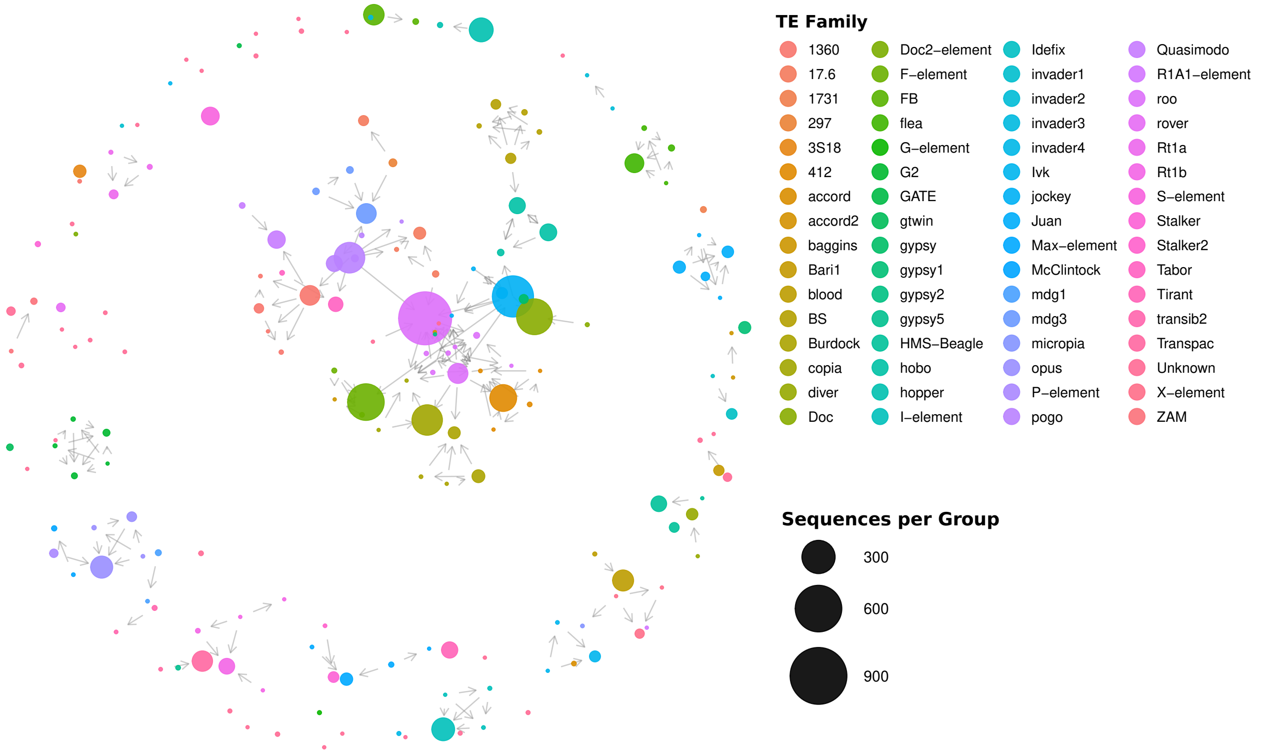


**Figure S14. Connection graph between TE insertions based on sequence similarity, color coded by TE family**. Nodes represent clusters of TEs with similar sequences; node size is proportional to cluster size. Most nodes (99.3%) are assigned to specific TE families. Directed edges denote one-way sequence similarity, indicating nesting of one TE (arrow origin) within another TE (arrow end).

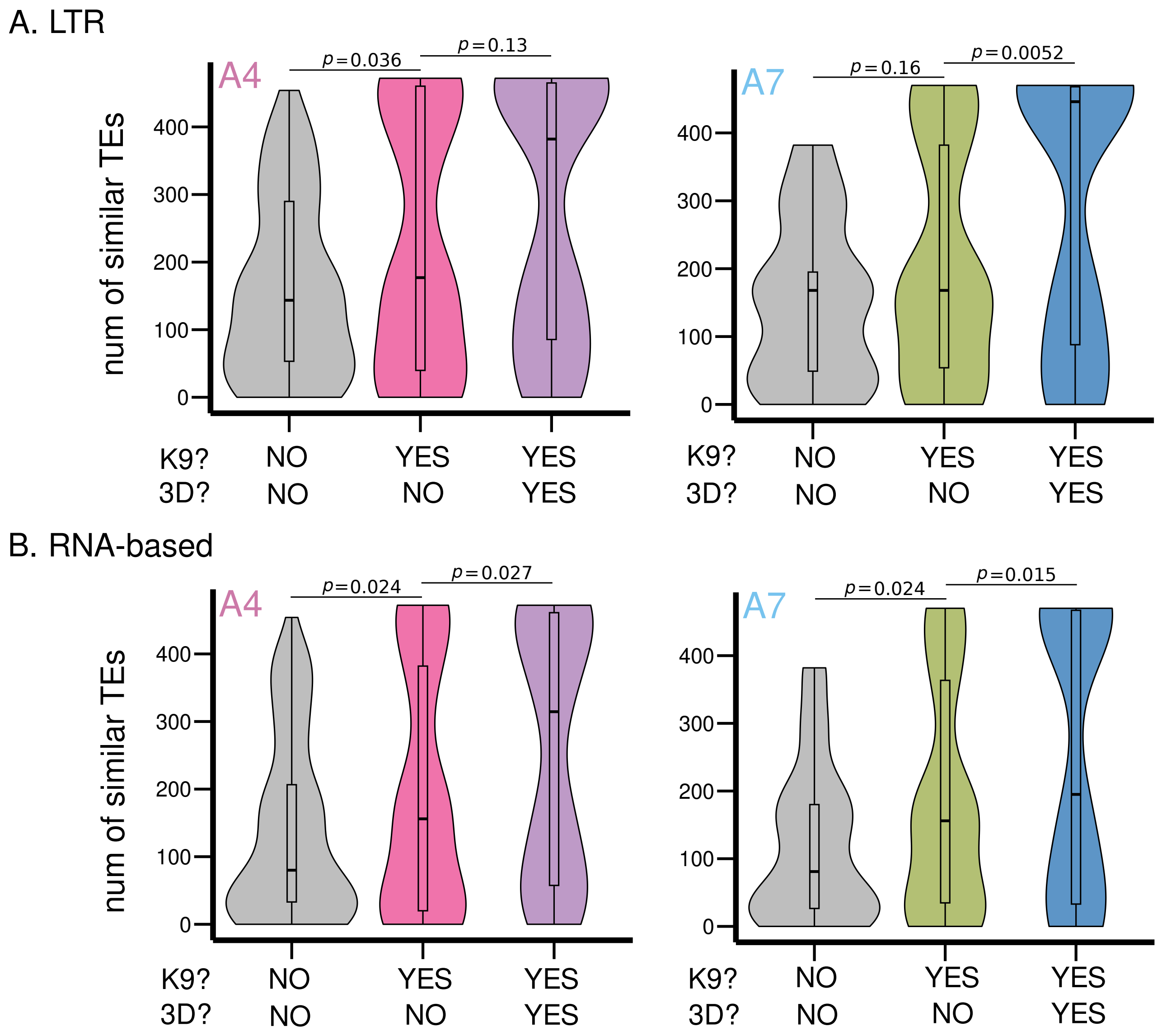


**Figure S15. Comparisons of number of similar TEs between categories of TEs restricting to certain TE type/class.** The number of similar TEs was compared between three categories of TEs: (1) without enrichment of H3K9me3 nor PCH 3D interactions, (2) H3K9me3-enriched TEs without PCH 3D interactions, and (3) H3K9me3-enriched TEs with PCH 3D interactions. Results are consistent irrespective whether the comparison was conducted for all TEs (Figure 4D) or a certain TE type: LTR (A) or class: RNA-based (B). Also see Table S2.

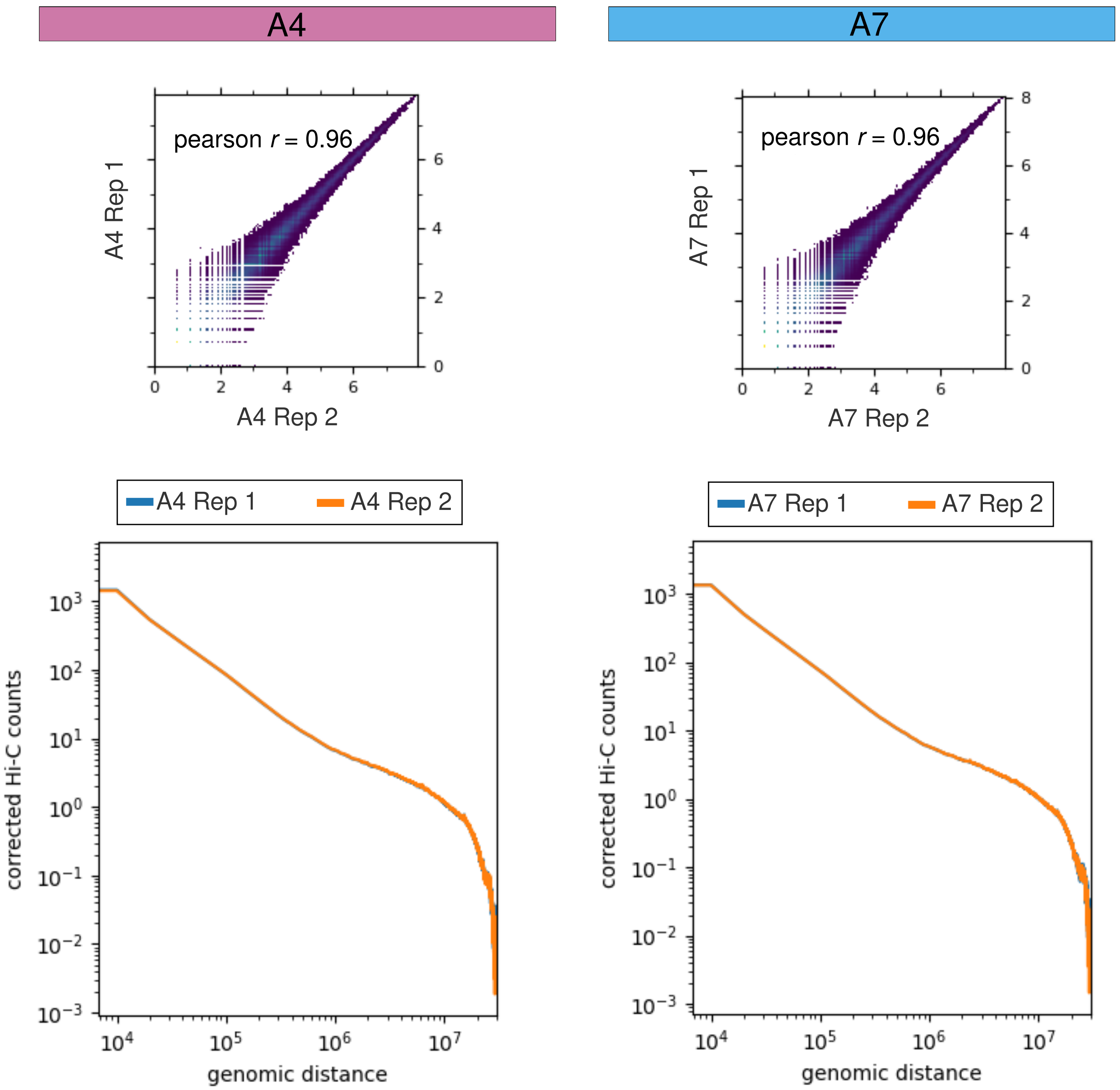


**Figure S16. Reproducibility of Hi-C biological replicates.** Top: Scatter plots comparing corrected Hi-C counts binned at 10kb between biological replicates for A4 and A7. Pearson correlation coefficients (A4: $r$ = 0.96, $p < 10^{-16}$; A7: $r$= 0.96, $p < 10^{-16}$) indicate high reproducibility. Bottom: Distance-dependent decay curves (log-log) for two replicates overlap nearly perfectly for both A4 and A7, indicating high reproducibility between biological replicates.

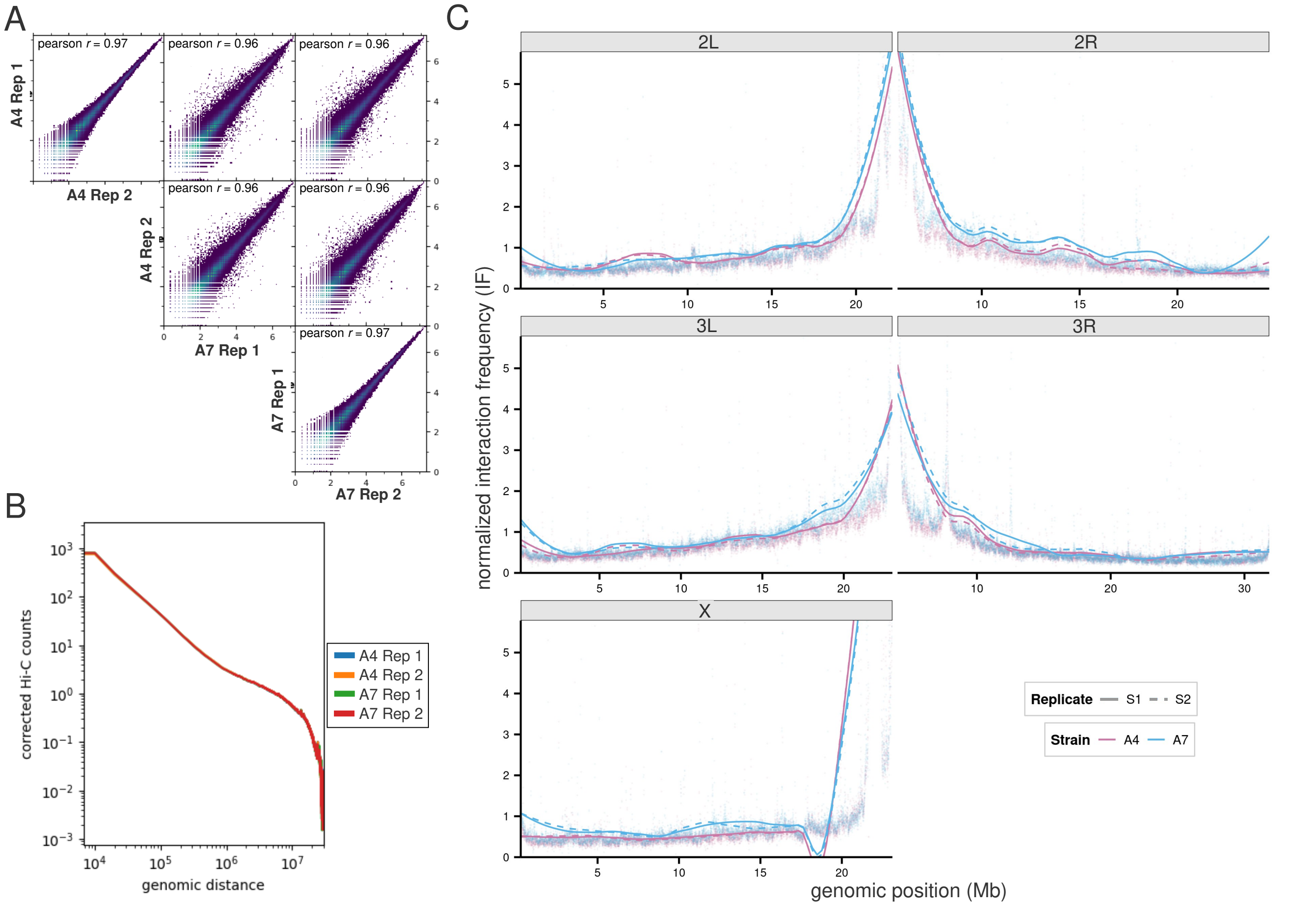


**Figure S17. Analysis of systematic biases between samples by aligning reads to the reference genome (A)** Scatter plots comparing Hi-C contact counts for A4 and A7 strains when mapped to the standard *D. melanogaster* iso-1 reference genome. The correlations between biological replicates (*Pearson correlation coefficient r* = 0.97) and between strains (*Pearson correlation coefficient r* = 0.96) are significantly high ($p < 10^{-16}$). **(B)** Distance-dependent decay curves (log-log) for A4 and A7 Hi-C samples aligned to the iso-1 reference, instead of their respective PacBio Hi-F assemblies. Curves overlap nearly perfectly, suggesting minimal differences. **(C)** Normalized interaction frequency (IF) along chromosome arms inferred from alignment of Hi-C data to iso-1 reference. A7 replicates (blue) consistently exhibit slightly higher interaction frequencies to PCH than A4 (pink). Such a result indicates that the differences we observed when aligning Hi-C data to strain-specific assemblies are not due to varying quality of genome assemblies, but likely the result of varying library complexity. Furthermore, these results underscore that even when replicates across strains appear globally similar, minor differences in library complexity can introduce systematic bias into comparative analyses, necessitating the correction strategy employed here.
